# Evolutionarily acquired activity-dependent transformation of the CaMKII holoenzyme

**DOI:** 10.1101/2023.01.10.523378

**Authors:** Shotaro Tsujioka, Ayumi Sumino, Yutaro Nagasawa, Takashi Sumikama, Holger Flechsig, Leonardo Puppulin, Takuya Tomita, Yudai Baba, Takahiro Kakuta, Tomoki Ogoshi, Kenichi Umeda, Noriyuki Kodera, Hideji Murakoshi, Mikihiro Shibata

## Abstract

Ca^2+^/calmodulin-dependent protein kinase II (CaMKII) has long been central in synaptic plasticity research. CaMKII is a dodecameric serine/threonine kinase that has been essentially conserved across metazoans for over a million years. While the mechanisms of CaMKII activation are well studied, its “behavior” at the molecular level has remained unobserved. Here, high-speed atomic force microscopy was used to visualize the activity-dependent structural dynamics of rat/hydra/*C. elegans* CaMKII in various states at nanometer resolution. Among the species, rat CaMKII underwent internal kinase domain aggregation in an activity-dependent manner and showed a higher tolerance to dephosphorylation by phosphatase. Our findings suggest that mammalian CaMKII has evolutionarily acquired a new structural form and a tolerance to phosphatase to maintain robust CaMKII activity for proper neuronal function.

**One-Sentence Summary:** High-speed atomic force microscopy reveals the activity-dependent structural dynamics of rat/hydra/*C. elegans* CaMKII

## Main Text

Ca^2+^/calmodulin-dependent protein kinase II (CaMKII) is a serine/threonine kinase with essential roles in various neuronal cell functions, such as long-term potentiation (LTP), long-term depression (LTD), learning and memory (*1–5*). Among four mammalian CaMKII isoforms (α, β, γ, and δ), α is the most abundantly expressed in the forebrain (*6, 7*). Notably, in the hippocampus, CaMKIIα represents ~2% of total proteins by mass (*8*) and is further concentrated (~7%) in the postsynaptic density (PSD) fraction (*9*).

CaMKIIα consists of four components: a kinase domain, a regulatory segment with a Ca^2+^/calmodulin (CaM)-binding site and three major phosphorylation sites (Thr286, Thr305, and Thr306), a linker region (residues 315–344), and a hub domain (Fig. 1A) (*4*). The structural features of the kinase domain, the regulatory segment, and the hub domain are well conserved across metazoans such as rat, hydra, and *Caenorhabditis elegans* (*C. elegans*) (fig. S1A). In contrast, the linker region is relatively variable among these three species, implying that the linker is possibly involved in distinct activation mechanisms and functions among species.

**Fig. 1.**
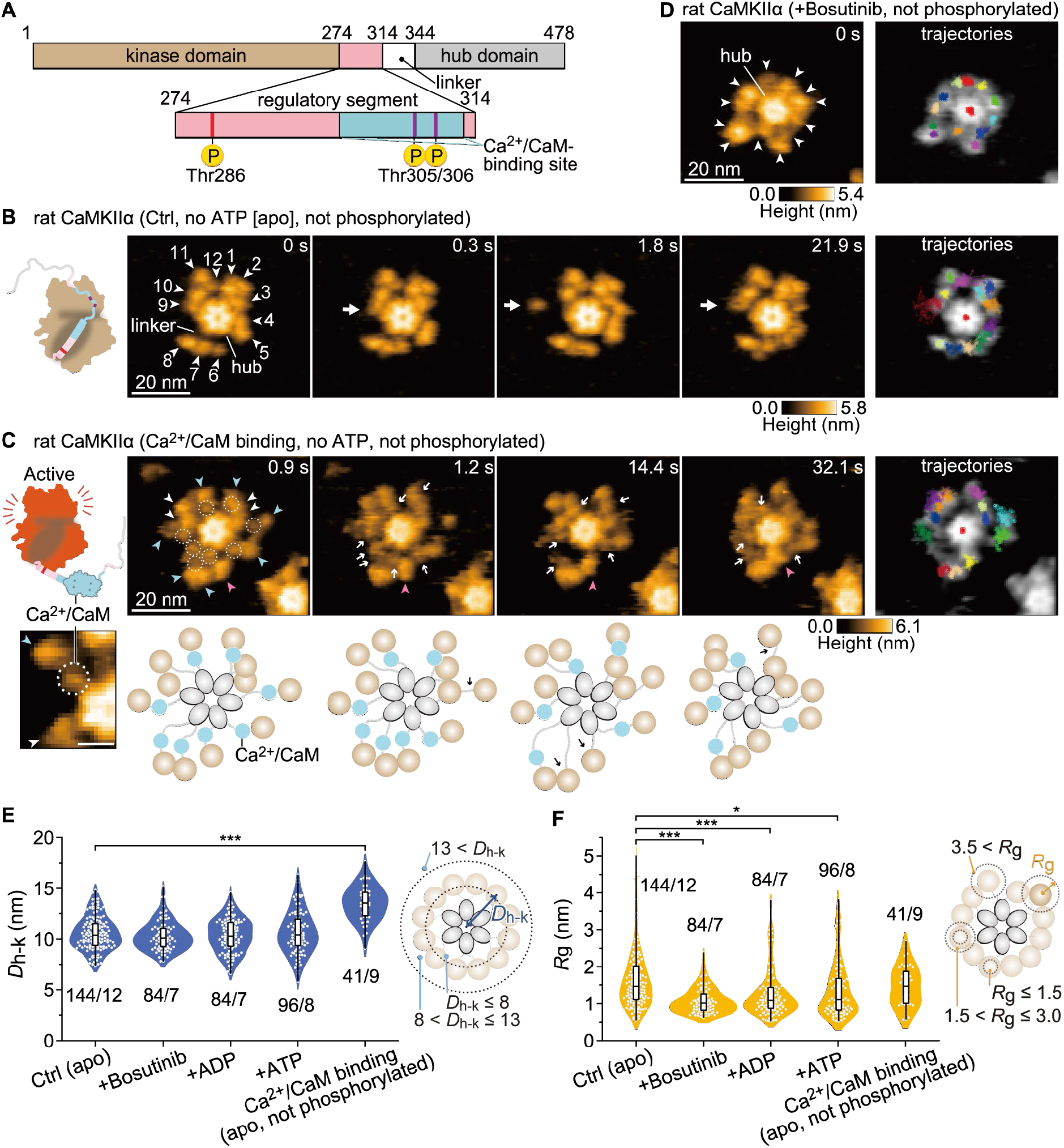
Ca^2+^/CaM binding causes extension of the CaMKIIα holoenzyme. (**A**) Domain structure of rat CaMKIIα. Numbers correspond to the amino acid sequence. (**B**) Sequential HS-AFM images of apo rat CaMKIIα (Ctrl, see also movie S1A). White arrowheads indicate kinase domains with arbitrary numbers. White arrows indicate the kinase domains in motion. Frame rate, 3.3 frames/s (also in C and D). Trajectories of the kinase domains and the center of the hub assembly (red in the center) were tracked for ~30 s (also in C and D). (**C**) Sequential HS-AFM images of Ca^2+^/CaM-bound rat CaMKIIα (1 mM Ca^2+^, 800 nM CaM, see also movie S2B). The dotted circles and white arrows indicate Ca^2+^/CaM. Blue and magenta arrowheads indicate Ca^2+^/CaM bound and dissociating kinase domains, respectively. Diagrams of the presumed structures of the linker regions in 12meric CaMKIIα are also shown at the bottom of each HS-AFM image. (**D**) HS-AFM images of rat CaMKIIα in the presence of 50 μM bosutinib (see also movie S2A). White arrowheads indicate kinase domains. (**E** and **F**) *D*_h-k_ (E) and *R*_g_ (F) for the respective conditions. The number of samples (kinases/holoenzymes) is indicated in the figure. (right) Illustrations of *D*_h-k_ and *R*_g_. \**p* < 0.05, \*\*\**p* < 0.001 (Kruskal–Wallis test with Dunnett’s post hoc test). HS-AFM experiments were repeated at least three times independently with similar results.

Mammalian CaMKIIα forms a double-ring structure consisting of 12 subunits connected via a hub domain (fig. S2A) (*10, 11*). Recently, single-particle electron microscopy showed that the CaMKIIα holoenzyme in the basal state adopts an extended conformation (*12*). In addition, while the kinase domain positioning is highly diverse, each kinase domain is autoinhibited by its regulatory segment (fig. S2B). This autoinhibition is released by the binding of activated CaM (Ca^2+^/CaM) to the regulatory segment, leading to kinase activation (fig. S2B). The adjacent activated kinase domains autophosphorylate each other at Thr286 (pT286). Even after Ca^2+^/CaM dissociation, the CaMKIIα subunit maintains pT286 and functions in adjacent kinase phosphorylation. This CaMKIIα autonomy has been hypothesized to be a form of molecular memory (as autonomy enables CaMKIIα to “remember” past Ca^2+^ stimuli by sustained activity) (*1–5*).

Upon pT286 and subsequent Ca^2+^/CaM dissociation, the secondary phosphorylation sites (Thr305/306) are exposed and autophosphorylated by both inter- (*trans*) and intramolecular (*cis*) subunits (fig. S2B) (*13–15*). Phosphorylation at Thr305/306 (pT305/306) inhibits subsequent Ca^2+^/CaM binding (*16*) and decreases the affinity for PSD (*17, 18*). These characteristics are associated with synaptic plasticity, such as LTP and LTD (*15, 19*).

While accumulated evidence has revealed the activation mechanisms and functions of CaMKII in synaptic plasticity, no one has previously observed the structural dynamics of CaMKII in liquid at the molecular level. Here, using high-speed atomic force microscopy (HS-AFM) (*20–22*), we visualized the activity-dependent structural dynamics of CaMKII in various states and in different species, including rats, hydras and *C. elegans*, in real time at nanometer resolution.

### The kinase domains in the CaMKIIα holoenzyme move freely around the hub assembly

For HS-AFM observation, CaMKIIα holoenzymes in the absence of ADP/ATP were purified from HEK-293 cells with two-step purification using His- and Strep-tags (fig. S3). Their activity was confirmed by western blotting (fig. S4). The purified proteins were immobilized on a positively charged mica surface modified with pillar[5]arene (*23*). This procedure enabled the visualization of single-particle dynamics of the CaMKIIα holoenzyme at high spatiotemporal resolution (<1 nm, 0.3 s/frame). In HS-AFM videos, individual particles appeared to consist of 12-mers with a central hub assembly and peripheral kinase domains in motion (more than 97% were 12-mer, and 3% were 14-mer [*n* = 96]) (Fig. 1B-D and movie S1A). To analyze the motion of kinase domains, the position of individual kinase domains in HS-AFM images was traced (Materials and Methods). Principal component analysis (PCA), which detected the interlocking motion of the 12 kinase domains, revealed that the main movement was circumferential around the central hub assembly (fig. S5). To quantify the mobility of each individual kinase domain, we determined the distance between the kinase domain and the center of the hub assembly (*D*_h-k_) and the radius of gyration *R_g_* (Materials and Methods). The results showed that each kinase domain was located at an average distance of 10.57 ± 0.36 nm from the center of the hub assembly with a large distribution of *R*_g_ (1.67 ± 0.25 nm [± SD], Ctrl in Fig. 1E and 1F), consistent with a previous study by electron microscopy (*12*). HS-AFM movie and analysis clearly showed that individual kinase domains are highly mobile.

Previous studies have shown that the linker length between the regulatory segment and the hub domain determines the size of the CaMKII holoenzyme (*24*).Consistent with this, our HS-AFM observations of the no-linker rat CaMKIIα (nlCaMKIIα) (lacking amino acids 315–344, fig. S1A) and its I321E (corresponding to I351E in the full-length rat sequence) mutant showed that both constructs take a more compact form than the wild type (fig. S6, movie S1). The distribution of *R*_g_ of nlCaMKIIα is smaller than that of the wild type, most likely due to the interactions of the kinase domains with the hub domain. Conversely, the distribution of *R*_g_ was increased by the introduction of I321E (fig. S6D), supporting the results of a previous report (*24*).

Next, we observed the CaMKII holoenzyme in the presence of bosutinib (SKI-606), an ATP-competitive inhibitor (*25*). Bosutinib was originally developed as an inhibitor of protein tyrosine kinases (*26*), and it was later found that it also binds to CaMKII in an ATP-competitive manner (*24, 27*). In the presence of bosutinib, *D*_h-k_ was comparable to that of apo CaMKIIα (Ctrl), consistent with a previous report showing that the size of the nlCaMKIIα holoenzyme is not altered by bosutinib binding (*24*). However, kinase motion (*R*_g_) was more restricted than in Ctrl (Fig. 1D-F and movie S2B): the binding of bosutinib to the ATP pocket most likely stabilizes the kinase domains and enhances the affinity with the regulatory segment. Similar results were obtained with ADP and ATP (Fig. 1E and 1F).

To further investigate, we monitored Ca^2+^/CaM binding to CaMKIIα by measuring Förster resonance energy transfer (FRET) by 2-photon fluorescence lifetime imaging microscopy (2pFLIM) (*28, 29*). mEGFP-fused CaMKIIα and mCherry-fused CaM were cotransfected into HeLa cells, and calcium ion influx was induced by ionophore bath application to activate CaM in the absence or presence of bosutinib (fig. S7). The results showed that bosutinib significantly reduced the efficiency of FRET, suggesting that Ca^2+^/CaM binding to CaMKIIα was predominantly inhibited (fig. S7B and S7C). Therefore, we speculate that the inhibition of Ca^2+^/CaM binding is due to the limited accessibility caused by increased interaction between the kinase domain and the regulatory segment.

### Ca^2+^/CaM binding induces the extended form of the CaMKIIα holoenzyme and the restricted motion of kinase domains

To visualize the Ca^2+^/CaM-bound CaMKIIα holoenzymes, we preincubated them for 5–20 min before performing HS-AFM observations. We avoided Ca^2+^/CaM incubation during HS-AFM observation because it largely increases the background noise due to CaM binding to the mica surface.

First, Ca^2+^/CaM-bound CaMKIIα was visualized in the absence of ATP (i.e., with no phosphorylation). It exhibited an extended form, estimated to be 2.94 nm more extended than that of Ctrl (Fig. 1C and 1E, movie S2C), probably due to the dissociation of the regulatory segment from the kinase domain. Small spherical objects, most likely Ca^2+^/CaM, were observed between the hub and kinase domains (dotted circles in Fig. 1C, movie S2C). HS-AFM also captured Ca^2+^/CaM dissociation and the concomitant approach of the kinase domain to the hub assembly (magenta arrows in Fig. 1C and fig. S8).

The average *R*_g_ (1.47 ± 0.54 nm) of Ca^2+^/CaM-bound CaMKIIα was comparable to that of Ctrl (Fig. 1F), but the highly mobile fraction (*R*_g_ greater than 3 nm) seen in Ctrl disappeared (Fig. 1F), suggesting that the mobility of kinase domains with Ca^2+^/CaM binding was restricted and that the kinase domains lay in an orderly arrangement. The restricted and orderly arrangement of kinase domains may facilitate efficient *trans*-phosphorylation in the oligomer.

### Ca^2+^/CaM-bound CaMKIIα holoenzymes with pT286 exhibit the extended form and have highly mobile kinase domains

Next, we preincubated CaMKIIα and Ca^2+^/CaM with ATP to prepare phosphorylated (pT286) CaMKIIα (fig. S4) and observed it under HS-AFM (Fig. 2A–C, movie S3). In CaMKIIα (Ca^2+^/CaM, pT286), Ca^2+^/CaM appeared between the hub and the kinase domains (dotted circles in Fig. 2A, movie S3B). Compared to that of nonphosphorylated CaMKIIα, the binding of Ca^2+^/CaM was more stable (the fractions of Ca^2+^/CaM bound longer than 60 s were 77.3% (*n* = 66) and 41.1% (*n* = 56) for pT286 and the nonphosphorylated state, respectively). This result is consistent with the CaM-trapping model (*30*). In addition, similar to CaMKIIα (Ca^2+^/CaM) (Fig. 2D), CaMKIIα (Ca^2+^/CaM, pT286) showed a structure extended in the radial direction by 3.3 nm. One of the remarkable differences is the increased *R*_g_ (Fig. 2E). We also imaged Ca^2+^/CaM-bound and Ca^2+^/CaM-unbound states of two different mutants (i.e., CaMKIIα_T286A_ and CaMKIIα_T305A/T306V_) (Fig. 2B–E, fig. S4 and S9, movies S4 and S5). We found that the structural change after Ca^2+^/CaM binding in the mutants was similar to that observed in the wild type.

**Fig. 2.**
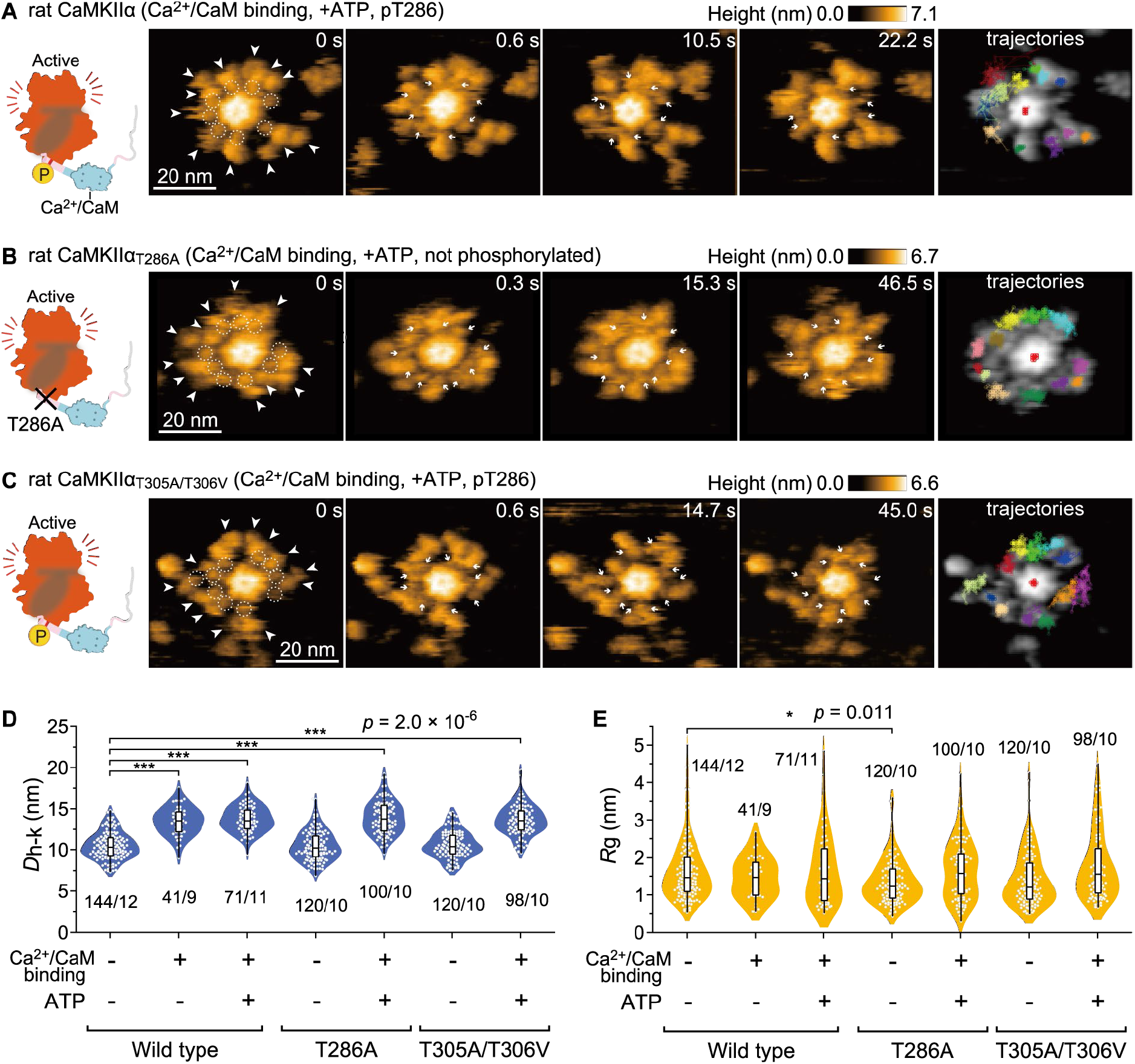
The CaMKIIα holoenzyme with pT286 in the Ca^2+^/CaM-bound state retains its extended structure and high mobility. (**A**) Sequential HS-AFM images (3.3 frames/s) of Ca^2+^/CaM-bound rat CaMKIIα with ATP (see also movie S3). CaMKIIα was activated by Ca^2+^/CaM and ATP (1 mM Ca^2+^, 800 nM CaM, 1 mM ATP) (also in B and C). White arrowheads indicate kinase domains. Dotted white circles and white arrows indicate Ca^2+^/CaM (also in B and C). (**B**) Sequential HS-AFM images of Ca^2+^/CaM-bound rat CaMKIIα_T286A_ with ATP (see also movie S4). (**C**) Sequential HS-AFM images of Ca^2+^/CaM-bound rat CaMKIIα_T305A/T306V_ with ATP (see also movie S5). (**D** and **E**) Distances from the center of the hub assembly to kinase domains (*D*_h-k_) (D) and gyration radii *R*_g_ (E) for the respective condition. \**p* < 0.05, \*\*\**p* < 0.001 (Kruskal–Wallis test with Dunnett’s post hoc test). The number of samples (kinases/holoenzymes) is indicated in the figure. HS-AFM experiments were repeated at least three times independently with similar results.

### Fully phosphorylated rat CaMKIIα holoenzymes (pT286/pT305/pT306) exhibit internal kinase aggregation

During HS-AFM observation of CaMKIIα (Ca^2+^/CaM, pT286), we found that the kinase domains formed stable dimers upon Ca^2+^/CaM dissociation (magenta arrows in Fig. 3A and third molecule in movie S3B). To clarify the molecular mechanisms of the stable oligomerization of kinase domains, we prepared Ca^2+^/CaM-unbound and fully phosphorylated CaMKIIα (pT286/pT305/pT306 in fig. S2B). In preparation, we incubated CaMKIIα in the presence of Ca^2+^/CaM and ATP, inducing pT286. Subsequently, Ca^2+^/CaM was released by incubation with an excess of EGTA, causing *trans*-/*cis*-phosphorylation at pT305/pT306 (fig. S4 and Fig. 3). In the HS-AFM videos, no Ca^2+^/CaM was observed in the CaMKIIα holoenzyme, indicating that CaM was dissociated in this experimental condition (Fig. 3B–D, movie S3C).

**Fig. 3.**
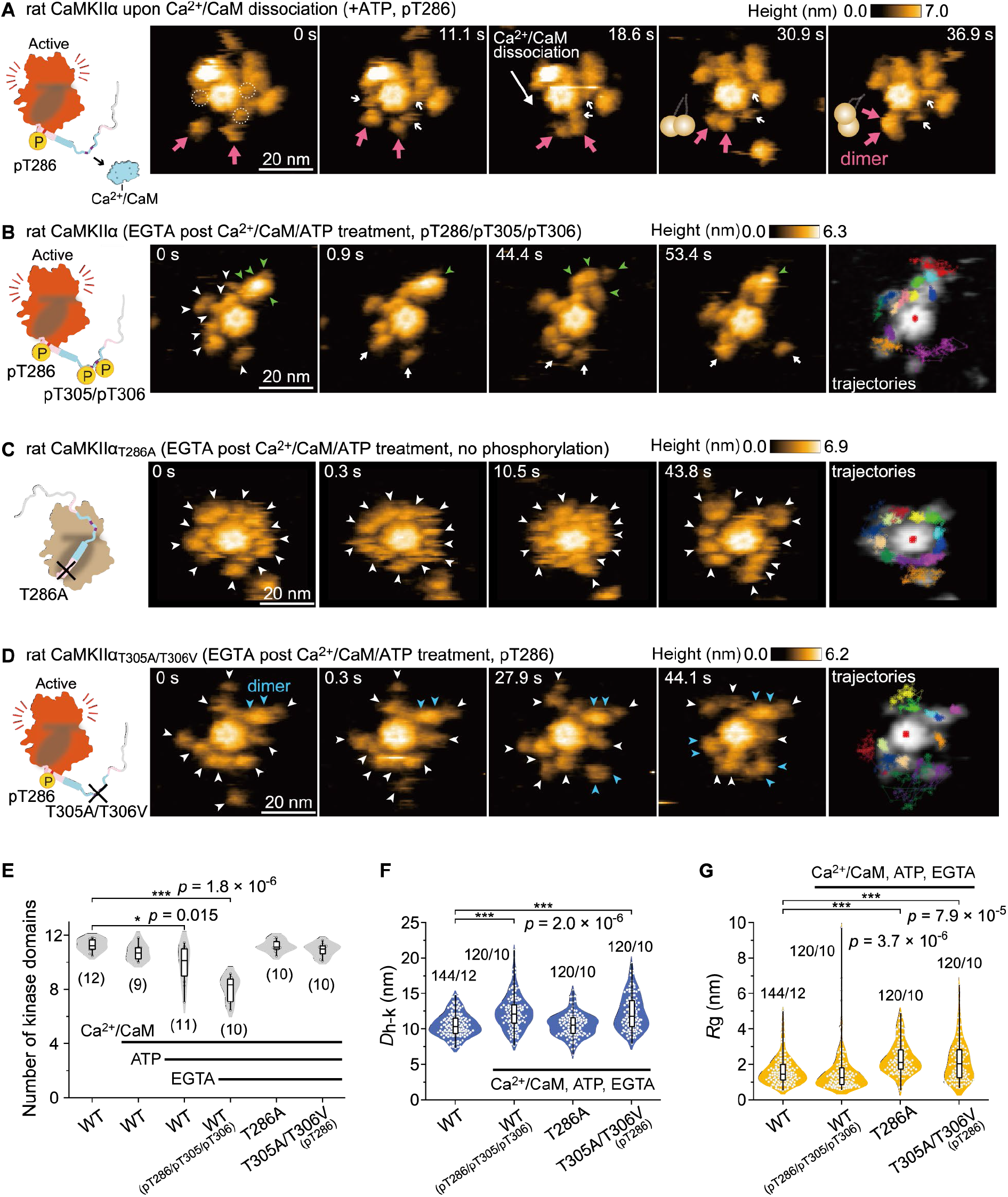
Fully phosphorylated rat CaMKIIα (pT286/pT305/pT306) exhibits kinase domain aggregation. (**A**) Sequential HS-AFM images of rat CaMKIIα at the time of stable dimer formation (see also movie S3B). CaMKIIα was activated by Ca^2+^/CaM and ATP (1 mM Ca^2+^, 800 nM CaM, 1 mM ATP). Dotted white circles and white arrows indicate Ca^2+^/CaM. Magenta arrows indicate the formation of stable dimers. Frame rate, 3.3 frames/s (also in B, C, and D). (**B**) Sequential HS-AFM images of pT286/pT305/pT306 rat CaMKIIα. CaMKIIα was first activated to induce pT286, as described in (A). Subsequently, EGTA (2 mM) was added to induce Ca^2+^/CaM dissociation and pT305/pT306 (autophosphorylation) (also in C and D). White, green, and blue arrowheads indicate kinase domains out, in aggregation, and in a dimer, respectively (also in C and D). (**C**) Sequential HS-AFM images of rat CaMKIIα_T286A_ (see also movie S4). Note that this mutant does not autophosphorylate at Thr286/Thr305/Thr306 (fig. S4, lanes #3–5). (**D**) Sequential HS-AFM images of pT286 rat CaMKIIα_T305A/T306V_ (see also movie S5). (**E**) Number of detectable kinase domains as the 4 nm object (unaggregated kinase domain) surrounding the hub assembly of CaMKIIα. The number of samples (holoenzymes) is indicated in the figure. (**F** and **G**) *D*_h-k_ (F) and *R*_g_ (G) for the respective conditions. \**p* < 0.05, \*\*\**p* < 0.001 (Kruskal–Wallis test with Dunnett’s post hoc test). The number of samples (kinases/holoenzymes) is indicated in the figure. HS-AFM experiments were repeated at least three times independently with similar results.

HS-AFM videos identified two states of kinase domains: aggregated and moving states (green and white arrows in Fig. 3B, movie S3C). To quantify the aggregated kinases, we determined the number of unaggregated kinase domains and found that the number significantly decreased in fully phosphorylated CaMKIIα (Fig. 3E).

The kinase domains (pT286/pT305/pT306) showed larger *D*_h-k_ (Fig. 3F) and broader distribution of *R*_g_ compared with Ctrl (Fig. 3G). The broader distribution is due to the coexistence of highly mobile unaggregated kinases and less mobile aggregated and stable kinases. As a further control experiment, we added ADP instead of ATP, and we confirmed that neither structural change nor aggregation occurred (fig. S10).

We next investigated the dependency of pT286 and pT305/pT306 on aggregation using CaMKIIα_T286A_ and CaMKIIα_T305A/T306V_. The results showed that CaMKIIαT286A did not exhibit aggregation (Fig. 3C and 3E, movie S4C). Note that this mutant showed no phosphorylation at Thr286/Thr305/Thr306 (fig. S4). Additionally, CaMKIIα_T305A/T306V_ exhibited no aggregation (Fig. 3D and 3E, movie S5C), although adjacent subunits temporarily formed a transient dimer (blue arrowheads in Fig. 3D).

In contrast to CaMKIIα_T286A_, CaMKIIα_T305A/T306V_ had a longer *D*_h-k_ (12.32 ± 2.76 nm; ± SD, Fig. 3F) and a larger *R*_g_ (2.50 ± 0.36 nm; ± SD, Fig. 3G), supporting the idea that pT286 causes an extended and highly mobile state. In summary, phosphorylation at both Thr286 and Thr305/306 is required for kinase aggregation.

### Hydra CaMKIIα and *C. elegans* CaMKII do not exhibit kinase aggregation in the fully phosphorylated state

CaMKII is highly conserved across species such as hydra and *C. elegans* (fig. S1A) (*24*). However, are the features of the structural dynamics seen in rat CaMKIIα also observable in other species? To answer this question, we applied HS-AFM to hydra CaMKIIα and *C. elegans* CaMKII, both of which are representative of the early stages of animal evolution. Similar to rat CaMKIIα, the kinase domains of both species moved freely around the central hub assembly in both the inactive and active (Ca^2+^/CaM, pT286) states (Fig. 4A–F, movies S6 and S7). In addition, they showed an extended structure upon Ca^2+^/CaM binding (Fig. 4B, 4E, and 4G).

**Fig. 4.**
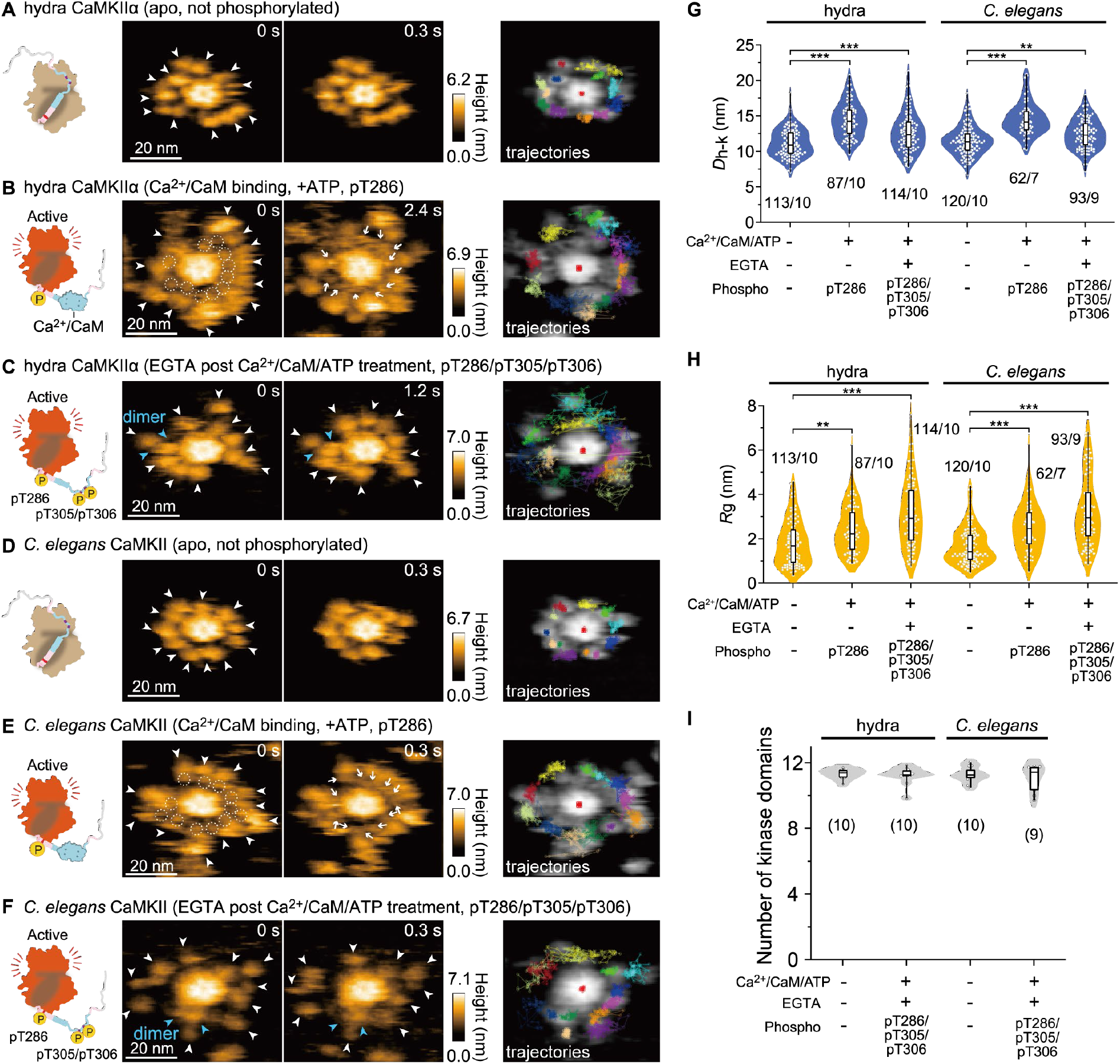
Hydra CaMKIIα and *C. elegans* CaMKII do not exhibit kinase aggregation. (**A** to **F**) HS-AFM images of hydra and *C. elegans* CaMKII (see also movies S5 and S6). The experimental conditions are described in the figures. Frame rate, 3.3 frames/s. In Figs. 4B and 4E, Ca^2+^/CaM-bound pT286 CaMKII was prepared by activated CaM and ATP (1 mM Ca^2+^, 800 nM CaM, 1 mM ATP). In Figs 4C and 4F, pT286 CaMKII was further treated with 2 mM EGTA to induce Ca^2+^/CaM dissociation and pT305/pT306 (autophosphorylation). White and blue arrowheads indicate kinase domains and in the dimer, respectively. (**G** and **H**) *D*_h-k_ (G) and *R*_g_ (H) for the respective conditions. The number of samples (kinases/holoenzymes) is indicated in the figure. \*\**p* < 0.01, \*\*\**p* < 0.001 (Kruskal–Wallis test with Dunnett’s post hoc test). HS-AFM experiments were repeated at least three times independently with similar results (also in I). **(I)** Number of detectable kinase domains as the 4 nm object (unaggregated kinase domain) surrounding the hub assembly of CaMKII. The number of samples (holoenzymes) is indicated in the figure.

Next, we observed hydra CaMKIIα and *C. elegans* CaMKII in the fully phosphorylated state (pT286/pT305/pT306) (fig. S4) and found that those kinase domains were not aggregated but highly mobile (Fig. 4C, 4F, 4H, and 4I). These data suggest that kinase aggregation in CaMKIIα at pT286/pT305/pT306 is mammalian specific.

To determine the biological function of this kinase aggregation, we examined whether kinase aggregation in rat CaMKIIα (pT286/pT305/pT306) alters its kinase activity. To this end, we assessed the kinase activity with a CaMKIIα substrate peptide, syntide-2 (*31*), by western blotting. Rat CaMKIIα (pT286/pT305/pT306), which is responsible for the kinase aggregation, more efficiently phosphorylated the substrate peptide (lanes #1/2 in fig. S11). A similar result was obtained with CaMKIIα_T305A/T306V_, which does not form kinase aggregates, suggesting that the increased kinase activity is not due to kinase aggregation (lanes #3/4 in fig. S11). However, we found that the kinase activity of hydra CaMKIIα was significantly reduced and that of *C. elegans* CaMKII was not increased in the fully phosphorylated state (fig. S11). This implies that structural differences between pT286 and pT286/pT305/pT306 alter the substrate specificity in hydra CaMKIIα and *C. elegans* CaMKII.

Next, we asked if kinase aggregation in rat CaMKIIα (pT286/pT305/pT306) leads to resistance to dephosphorylation. To assess this hypothesis, we incubated phosphorylated CaMKIIα with PP2A, a major CaMKII phosphatase (*17, 32*), and quantified phosphorylation at Thr286 by western blotting (fig. S12A, B). Phosphorylated rat CaMKIIα (wt) was dephosphorylated in a PP2A concentration-dependent manner, and similar results were obtained for the nonaggregate CaMKIIα mutant (T305A/T306V) (fig. S12C). This result suggests that kinase aggregation does not increase tolerance to PP2A compared with the nonaggregated form in rat CaMKIIα. In contrast, we found that rat CaMKIIα is more resistant to PP2A than hydra CaMKIIα in a broad concentration range (fig. S12D). Rat CaMKIIα was also more resistant than *C. elegans* CaMKII at high PP2A concentrations (fig. S12E).

## Discussion

CaMKII has long been studied using various methods, such as biochemical assays and electron microscopy. However, single-molecule CaMKII holoenzymes have never been visualized in action. Here, using HS-AFM, we observed the activity-dependent structural dynamics of CaMKII from different species, including rats, hydras, and *C. elegans*. Built upon the analyses of our observations, we described (i) the effects of ADP/ATP/bosutinib binding, (ii) the stabilization of kinase domains upon Ca^2+^/CaM binding, (iii) the autophosphorylation-dependent kinase aggregation in mammalian CaMKIIα but not in older nonmammalian species, and (iv) the higher phosphatase tolerance of mammalian CaMKIIα than that of nonmammalian species. Based on our HS-AFM results, we propose an activation mechanism of CaMKII (Fig. 6).

Previously, 3D EM reconstruction with pseudoatomic resolution showed that the CaMKII holoenzyme in the basal state is in an extended conformation and that its kinase domain positioning is highly flexible (*12*). Consistent with this view, we found that the kinase domains in the CaMKIIα holoenzyme at the basal state are indeed in a mobile state, most likely facilitating Ca^2+^/CaM binding to CaMKIIα. The existence of kinase domain dimers was predicted (*12, 33*); however, in our observation, most domains existed as monomers. This discrepancy may be due to the methodological difference between the studies.

We assessed the rat CaMKIIα holoenzyme structure in the absence and presence of bosutinib. Consistent with a previous study (*24*), no size change in rat CaMKIIα was observed. However, our observations revealed that bosutinib binding restricts kinase domain motion. As observed by 2pFLIM, bosutinib inhibits Ca^2+^/CaM binding to CaMKIIα in HeLa cells. Bosutinib binding probably enhances the affinity between the kinase domain and the regulatory segment and inhibits Ca^2+^/CaM binding.

Upon Ca^2+^/CaM binding, the CaMKIIα holoenzymes extend by ~3 nm from the center of the hub assembly, probably dissociating the regulatory segment from the active site in the kinase domain and exposing the Thr286 phosphorylation site in the regulatory segment, as previously suggested (*34*). In addition, Ca^2+^/CaM binding unexpectedly decreases the kinase domain mobility, which then increases again with pT286, even with Ca^2+^/CaM binding. We speculate that Ca^2+^/CaM binding stabilizes the α-helical structure of the regulatory segment and increases the accessibility of the adjacent kinase domains to Thr286. Thr286 phosphorylation changes the structure of the regulatory segment from α-helical to disordered, and the mobility of the kinase domain increases. This pT286-dependent structural transition might be supported by previous results showing that pT286 dramatically increases the affinity to Ca^2+^/CaM (CaM-trap) (*30*).

Upon Ca^2+^/CaM dissociation, the regulatory segment with pT286 remains free from the active site and is *trans*-/*cis*-phosphorylated at the second phosphorylation site (Thr305/306) (*14, 15*). Using HS-AFM, we found that the rat CaMKIIα holoenzyme with pT286/pT305/pT306 phosphorylation exhibits internal kinase aggregation. We speculate that pT286/pT305/pT306 phosphorylation induces disorder in the regulatory segments, leading to a string-like entanglement structure and increasing the interregulatory segment interaction, as seen in the microtubule-associated protein tau (*35*).

Several possible physiological implications of kinase aggregation in rat CaMKIIα arise. First, the aggregation of kinase domains would affect kinase activity. Accordingly, the kinase aggregation observed here offers a new regulatory mechanism for kinase activity. Indeed, a kinase assay showed that the CaMKIIα holoenzyme with pT286/pT305/pT306 phosphorylates its substrate more efficiently. This result does not contradict a previous study showing Thr305/306-dependent substrate phosphorylation (*15*). However, the nonaggregate mutant (i.e., T305A/T306V) also showed increased kinase activity. Thus, the aggregation of kinase domains may not be the direct cause of increased kinase activity. Second, kinase aggregation might limit the accessibility of phosphatases to pT286/pT305/pT306 by hiding them in aggregates. However, since both aggregated and nonaggregated forms of CaMKIIα exhibited similar phosphatase tolerance, it is unlikely. Third, kinase aggregation might regulate the affinity of the CaMKIIα holoenzyme for binding partners by limiting its accessibility to kinase domains. For instance, previous studies have reported that pT305/pT306 reduces affinity with PSD in vitro (*17*), while the introduction of T305A/T306A increases association with PSD. This increased association lowers the threshold for hippocampal LTP and results in more rigid and less fine-tuned hippocampal-dependent learning (*19*). In addition, a recent study suggests that LTD stimulation phosphorylates at Thr305/306 over Thr286 and promotes CaMKII translocation into inhibitory synapses (*15*).

CaMKII is an evolutionarily conserved protein (*36*). However, its linker region is hypervariable among species (fig. S1A). To date, the influence of linker differences on CaMKII functions has not yet been investigated. This study revealed the structural and dynamic differences among rat, hydra, and *C. elegans* CaMKII holoenzymes, which were visualized in liquid by HS-AFM. In the basal state and Ca^2+^/CaM-bound pT286, all species of CaMKII holoenzymes exhibited similar *D*_h-k_ and *R*_g_. Previously, crystal structural analyses combined with SAXS measurements suggested that the *C. elegans* kinase domain forms dimers (*33*). However, our HS-AFM images showed that the *C. elegans*, rat and hydra kinase domains all exist as monomers in the basal state and as Ca^2+^/CaM-bound pT286. This discrepancy may be due to methodological differences between the studies. The main difference between CaMKIIα species was found in their pT286/pT305/pT306. In rat CaMKIIα, kinase domains exhibited aggregation, but ancestral CaMKIIα (i.e., hydra and *C. elegans* CaMKII) did not. The amino acid sequences of the regulatory segment and linker region differ among species (fig. S1A), which may explain the difference in kinase aggregation.

One of the remarkable findings in this study is the differential species-dependent phosphatase tolerance in pT286/pT305/pT306. Biochemical assays found that rat CaMKIIα is more resistant to PP2A than hydra CaMKIIα in a broad concentration range (fig. S12D). Rat CaMKIIα was also more resistant than *C. elegans* CaMKII at high PP2A concentrations (fig. S12E). The mechanisms of the higher tolerance of mammalian CaMKII compared to that of hydra CaMKIIα and *C. elegans* CaMKII are unknown. One possible cause of this difference is the difference in *R*_g_ (fig. S13). Since *R*_g_ in pT286/pT305/pT306 of nonmammalian CaMKII is larger than that of rat CaMKIIα, nonmammalian CaMKII may have a larger free space around the regulatory segment, which could facilitate phosphatase access to phosphorylated regulatory segments for dephosphorylation.

In conclusion, our findings provide the basis for a deeper understanding of the molecular mechanisms of CaMKIIα activation. In particular, mammalian CaMKIIα-specific structural arrangement and phosphatase tolerance may be critical features for Ca^2+^ signal integration and sustained activation of CaMKIIα required for LTP, LTD, learning, and memory. In addition, these evolutionarily acquired features may distinguish neuronal function between mammals and other species.

**Fig. 5.**
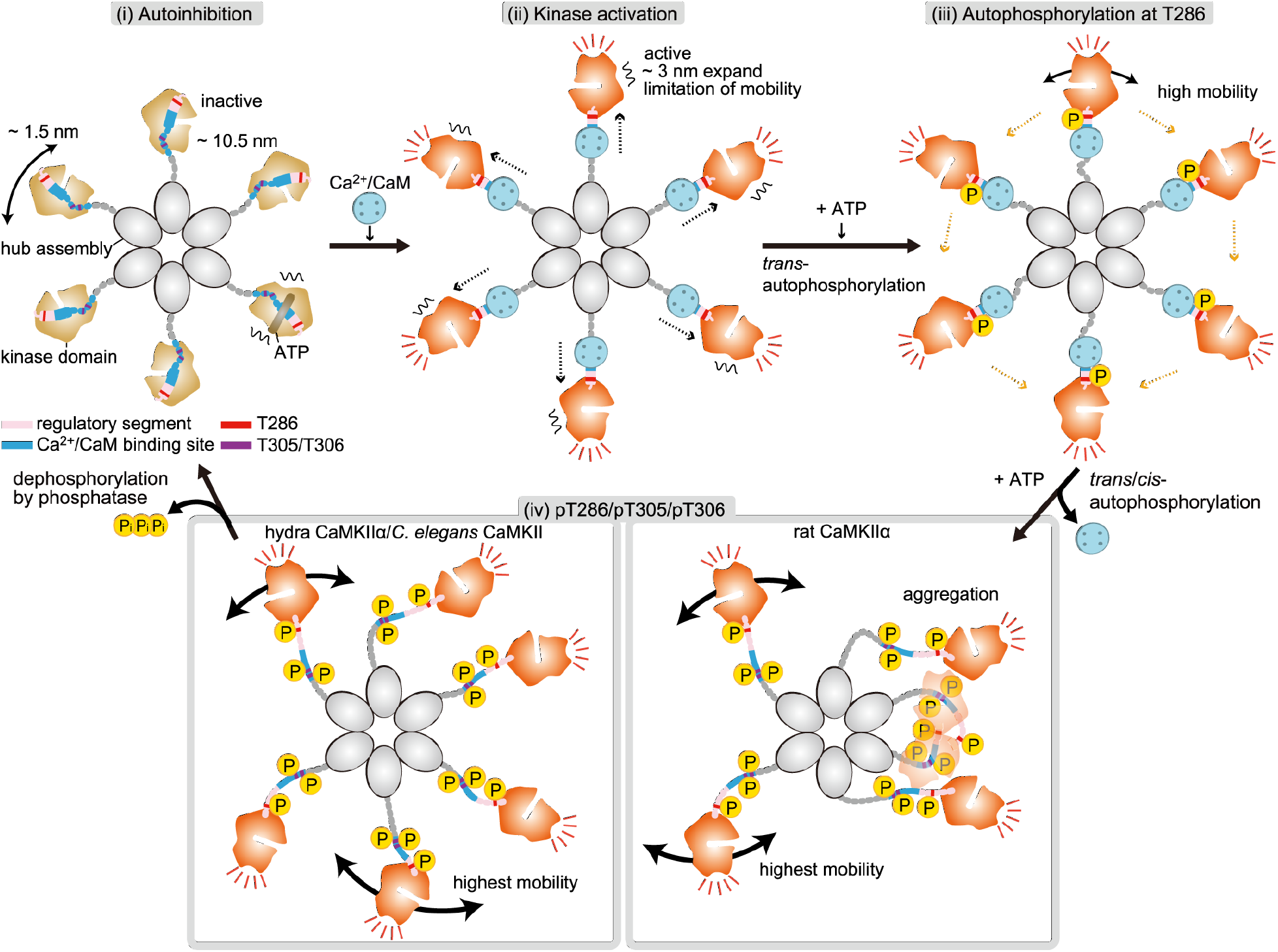
Schematic of CaMKIIα dynamics in the process of activation. A dodecameric CaMKIIα holoenzyme is depicted as a hexamer for simplicity. (i) In an inactive state, kinase domains are distributed within ~10.5 nm in radius from the center of the hub assembly with some mobility. (ii) Ca^2+^/CaM binding leads to the extended form of the holoenzyme and mobility reduction of kinase domains. (iii) Subsequent Thr286 autophosphorylation (autophosphorylation at T286) leads to higher kinase mobility. (iv) Dissociation of Ca^2+^/CaM induces an increase in kinase mobility (highest mobility) and causes autophosphorylation at Thr305/Thr306. The pT286/pT305/pT306 state of rat CaMKIIα exhibits kinase aggregation, while hydra CaMKIIα and *C. elegans* CaMKII do not. The phosphorylated state is maintained until dephosphorylation by phosphatase occurs.

## Acknowledgments

We thank Dr. Ryohei Yasuda at Max Planck Florida Institute and Dr. Toshio Ando at Kanazawa University for providing the HS-AFM apparatus, Dr. Takayuki Uchihashi at Nagoya University for providing HS-AFM software, and Yuka Kawahata and Yoko Mikami at Kanazawa University for collecting HS-AFM images and laboratory management. Calculations were partly conducted on a supercomputer at the Research Center for Computational Science in Okazaki, Japan.

## Author contributions

Conceptualization: HM, MS

Data curation: ST, AS, YN, TS, HF, HM, MS

Formal Analysis: AS, TS, HF, LP, MS

Funding acquisition: TO, HM, MS

Investigation: HM, MS

Methodology: KU, NK, MS

Project administration: HM, MS

Resources: TT, YB, TK, TO, HM, MS

Software: TS, HF, KU

Supervision: HM, MS

Validation: ST, MS

Visualization: AS, TS, HF, LP, HM, MS

Writing – original draft: HM, MS

Writing – review & editing: AS, TS, HF, LP, KU, NK, HM, MS

## Data and materials availability

Data will be available upon reasonable request.

## Supplementary Materials

Materials and Methods

Figs. S1 to S13

References (37–48)

Captions for Movies S1 to S7

Movies S1 to S7

