## Supplemental Figs for "Evolutionarily acquired activity-dependent transformation of the CaMKII holoenzyme"

Shotaro Tsujioka *et al.*

**This PDF file includes:**

Materials and Methods
Figs. S1 to S13
References (37–48)
Captions for Movies S1 to S7

**Other Supplementary Materials for this manuscript include the following:**

Movies S1 to S7

Reagent

A calcium ionophore (4-bromo A-23187) was purchased from Cayman Chemical (USA). Bosutinib (Cat. No. 4361) was purchased from Tocris (UK).

Plasmids

The rat (*Rattus norvegicus*) CaMKIIα gene encoding CaMKIIα (a.a. 1–478) was a gift from Y. Hayashi at Kyoto University. The synthesized gene encoding rat (*Rattus norvegicus*) calmodulin (CaM) was purchased from Eurofins (Tokyo, Japan). The synthesized genes encoding hydra (*Hydra vulgaris*) CaMKIIα isoform X2 and *C. elegans* (*Caenorhabditis elegans*) CaMKII were purchased from Genscript (Tokyo, Japan). For PP2A-related genes (*37*), human PPP2CA was purchased from Promega (WI, USA), and synthesized human PTPA (PPP2R4) and Mini-A (PPP2R1A) were purchased from FASMAC (Kanagawa, Japan).

The plasmid containing 6×His/Strep tags (MDYKDDDD<u>HHHHHH</u>K<u>WSHPQFEK</u>GTGGQ QMGRDLYDDDDKDLYKSGLRSRA) fused to the N-terminus of rat/hydra/*C. elegans* CaM KII/PPP2CA/PTPA/Mini-A was inserted into a modified pEGFP-C1 mammalian expression v ector (the kanamycin resistance gene was replaced with the ampicillin resistance gene) by r eplacing EGFP. To construct the rat no-linker CaMKIIα (nlCaMKIIα) plasmid, the sequence encoding a.a. 315–344 was removed from the rat CaMKIIα sequence. CaMKII mutants with point mutations were constructed using the QuikChange site-directed mutagenesis kit (Agile nt Technologies, USA). To construct 6×His-tagged CaM and sfGFP-fused syntide-2 peptide (amino acid sequence PLARTLSVAGLPGKK) plasmids, the respective genes were inserted i nto the pRSET bacterial expression vector (Invitrogen, USA). mEGFP-CaMKIIα and mCherr y-CaM were constructed by insertion into the modified pEGFP-C1 vector, replacing EGFP.

Protein purification from bacteria

The His-tagged CaM and sfGFP-fused Syntide-2 were overexpressed in *Escherichia coli* (DH5α), and the culture medium (~250 ml) was centrifuged at 2,500 ×g for 20 min. The cell pellet was dissolved in 10 ml of ice-cold phosphate-buffered saline (PBS) buffer with 1% Triton X-100 and 5 mM imidazole, sonicated, and then centrifuged at 20,000 × g for 10 min. The supernatant was loaded into a $Ni^+$-nitrilotriacetate (NTA) column (HiTrap, GE Healthcare, USA). The NTA column bound to the protein was washed with 5 mM and 50 mM imidazole containing 20 mM Tris-HCl pH 7.4 buffer and eluted with 20 mM Tris-HCl/500 mM imidazole buffer. The concentration of the purified protein was measured using the Bradford assay (Bio-Rad, U.S.A.) by comparing BSA as a standard.

Protein purification from HEK293 cells

All CaMKII proteins and PP2A were prepared with the following method. 6×His/Strep tagged CaMKIIα was expressed in HEK293 cells. Briefly, HEK293 cell culture was maintained on 15-cm plates in Dulbecco's modified Eagle's medium (DMEM) supplemented with 5% fetal bovine serum (FBS) with no antibiotics at 37°C and 5% $CO_2$. Three 15-cm dishes at 70% confluency were prepared for transfection. The cells were transfected with the plasmid encoding CaMKII genes using Lipofectamine 2000 following the manufacturer's protocol (Thermo Fisher Scientific, USA). For PP2A, three plasmids encoding PPP2CA/PTPA/Mini-A genes were cotransfected. One day after transfection, the cells were collected and lysed with 10 mL of ice-cold phosphate-buffered saline (PBS) buffer with 1% Triton X-100 and 5 mM imidazole and then centrifuged at 20,000 × g for 10 min. The supernatant was loaded into Strep-Tactin Sepharose (Nacalai, Japan), washed with 20 mM Tris-HCl/150 mM KCl, and subsequently eluted with 2.5 mM desthiobiotin, pH 7.4. The pooled proteins were further purified with an NTA column, and the concentration of CaMKII proteins was determined as described above. The PP2A concentration was determined as described in the following section. The purified proteins were stored in 20 mM Tris-HCl/150 mM KCl with 50% glycerol and 2 mM DTT at -30°C before experiments.

Measurement of purified PP2A activity

One unit is defined as the amount of PP2A that hydrolyzes 1 nmol of 50 mM p-nitrophenyl phosphate (pNPP) in 1 minute at RT in a total reaction volume of 100 µl. The phosphatase activity was assayed in 100 µl of the reaction mixture (50 mM pNPP, 50 mM Tris pH 7.6, 150 mM KCl, 10 mM $MgCl_2$, 1 mM $MnCl_2$). The reaction was initiated by the addition of 1 µl of PP2A. After 3 min, the amount of p-nitrophenol was determined by reading the absorbance at 405 nm (18,000 $M^{-1}$ $cm^{-1}$). This method gives PP2A activity in unit/µl, and our purified PP2A typically had activity of ~0.5 unit/µl.

Biochemical assay for phosphorylation/dephosphorylation of CaMKII

Standard kinase assays were performed at 30°C for 5 min incubation with 50 nM of purified CaMKII mutants, 800 nM CaM, 1 mM $CaCl_2$, and 1 mM ATP in reaction buffer (50 mM Tris-HCl, pH 7.4, 150 mM KCl, 10 mM $MgCl_2$). To release CaM from CaMKII, 2 mM EGTA was added and incubated for 5 min at 30°C and an additional 10 min at 25°C to match the conditions for AFM observation (i.e., after the sample was loaded on the AFM apparatus, it typically takes 10 min before observation). For substrate phosphorylation, sfGFP-Syn1 was added at 1 µM and incubated for 20 min at 25°C. For CaMKII dephosphorylation by PP2A, phosphorylated CaMKII proteins were incubated with the indicated concentration of PP2A for 10 min at 25°C. The reactions were stopped by adding SDS sample buffer and then analyzed by western blotting.

Western blotting was performed with the following antibodies: anti-phospho-CaMKII (Thr286) (D21E4; Cell Signaling Technology, USA); anti-pT305 (abx012403; abbexa); anti-phospho-CaMKII (Thr306) (p1005-306; PhosphoSolutions, USA); anti-phospho-PKA substrate (RRXS*/T*) (#9624; Cell Signaling Technology); anti-His-tag (27E8; Cell Signaling Technology, USA); and HRP-anti-rabbit/mouse (Jackson Laboratory, USA).

2-photon FLIM-FRET experiment

HeLa cells were cultured in DMEM supplemented with 5% fetal bovine serum at 37°C and 5% $CO_2$. The cells were transfected with the plasmids (i.e., mEGFP-CaMKIIα and mCherry-CaM) by means of Avalanche-Everyday transfection reagent (EZ biosystems, USA), followed by incubation for 16–20 h. FLIM-FRET imaging was conducted in a solution containing 4-(2-hydroxyethyl)-1-piperazineethanesulfonic acid (HEPES; 30 mM, pH 7.3)-buffered artificial cerebrospinal fluid (130 mM NaCl, 2.5 mM KCl, 1 mM $CaCl_2$, 1 mM $MgCl_2$, 1.25 mM $NaH_2PO_4$, 25 mM glucose) at room temperature (23–25°C).

Details of the 2-photon FLIM-FRET imaging are described elsewhere (29, *38*). Briefly, mEGFP-CaMKIIα was excited with a Ti:sapphire laser (Mai Tai; Spectra-Physics, U.S.A.) tuned to 920 nm. The scanning mirror was controlled with ScanImage software (*39*). The green fluorescence photon signals were collected by an objective lens (60×, 1.0 NA; Olympus, Japan) and detected by a photomultiplier tube (H7422-40p; Hamamatsu, Japan) placed after a dichroic mirror (565DCLP; Chroma Technology, USA) and emission filter (FF01-510/84; Semrock, USA). Measurement of fluorescence lifetime was processed using a time-correlated single-photon counting board (SPC-150; Becker & Hickl, Germany) controlled with custom software (*38*). For fluorescence lifetime image construction, the mean fluorescence lifetime in each pixel was translated into a color-coded image (*29*). Analysis of the lifetime change and binding-fraction change was conducted as described elsewhere (*29*).

HS-AFM apparatus

HS-AFM observations were performed using a homemade high-speed atomic force microscope (*22, 40-41*). An optical beam deflection detector was used to detect the cantilever (BL-AC10DS-A2, Olympus, Japan) deflection in tapping mode using an infrared (IR) laser at 780 nm and 0.7 mW. The IR beam was focused onto the back of the cantilever through a 50× objective lens (TU Plan Apo EPI 50X, Nikon, Japan). The reflection of the IR beam from the cantilever was detected with a two-segmented PIN photodiode (MPR-1, Graviton, Japan). The spring constant of the cantilever was ~100 pN/nm. The resonant frequency and quality factor of the cantilever in a liquid were ~400 kHz and ~2, respectively. Although the cantilever has the original bird-beak-like triangular portion as an AFM tip, to improve the spatial resolution of HS-AFM, an amorphous carbon tip was grown on the original AFM tip by electron beam deposition (EBD) using SEM. The length of the additional AFM tip was ~500 nm, and the apex of the tip was ~1 nm in radius after further plasma etching by a plasma cleaner (Tergeo, P.I.C. Scientific, USA). All HS-AFM images were obtained from cantilevers with additional AFM tips. The free oscillation amplitude of the cantilever was under 1 nm, and the set-point amplitude was set to 80% of the free amplitude during HS-AFM observations. To reduce the force between the sample and AFM tip, a recently developed "only trace imaging" (OTI) mode was used for HS-AFM scanning (42). HS-AFM data were collected using laboratory-developed software based on Visual Basic.NET (Microsoft).

Substrate for HS-AFM observations

For all HS-AFM observations, we modified a mica surface using cationic C2 pillar[5]arene (P[5]A+) to change the surface charge from negative to positive (43). The electrostatic potential map of CaMKIIα shows that the surface charge of the hub assembly is partially negative (12), while the surface charge of the kinase domains is partially positive. Indeed, the mobility of the kinase domains of CaMKIIα was strongly inhibited on negatively charged bare mica due to electrostatic interactions. To prevent the inhibition of kinase domain mobility on an HS-AFM substrate, we used P[5]A+ as an AFM substrate. An aqueous solution of 70 µM P[5]A+ was deposited onto a freshly cleaved mica substrate (1.0 mm in diameter, Furuuchi Chemical, Japan). P[5]A+ was incubated at room temperature on the mica surface for 15 min, and the surface was rinsed with Milli-Q water to remove unadsorbed P[5]A+.

HS-AFM observations

CaMKII holoenzymes in the basal state were observed in buffer A (50 mM Tris-HCl, pH 7.4, 15 mM KCl, 10 mM $MgCl_2$, 10% glycerol) with 0.1 mM EGTA. For the inhibitor experiment, 50 nM CaMKIIα was premixed at 30°C for 5 min with 50 µM bosutinib in buffer B (50 mM Tris-HCl, pH 7.4, 150 mM KCl, 10 mM $MgCl_2$) with 0.1 mM EGTA. Then, HS-AFM observations were performed in buffer A with 0.1 mM EGTA and 50 µM bosutinib. For $Ca^{2+}$/CaM-bound CaMKII holoenzymes, 50 nM CaMKII and 800 nM CaM were premixed in buffer B with 1 mM $CaCl_2$ and incubated at 30°C for 5 min. Then, observations were performed in buffer A with 1 mM $CaCl_2$.

For $Ca^{2+}$/CaM-bound CaMKII holoenzymes in the presence of ATP, we premixed 50 nM CaMKII and 800 nM CaM in buffer B with 1 mM $CaCl_2$ and 1 mM ATP and incubated the mixture at 30°C for 5 min. Then, HS-AFM was performed in buffer A with 1 mM ATP.

For CaMKII holoenzymes in EGTA and ATP, we premixed 50 nM CaMKII and 800 nM CaM in buffer B with 1 mM $CaCl_2$ and 1 mM ATP and incubated at 30°C for 5 min. After that, to dissociate $Ca^{2+}$/CaM from CaMKIIα, we added 2 mM EGTA and incubated at 30°C for an additional 5 min. Then, HS-AFM was performed in buffer A with 2 mM EGTA and 1 mM ATP. All HS-AFM experiments were performed at room temperature (24–26°C).

HS-AFM image processing and data analysis

HS-AFM images were processed using Fiji (ImageJ) software (NIH, USA) (44). A 0.5-pixel-radius mean filter was applied to each HS-AFM image to reduce noise. The Template Matching and Slice Alignment plugin for ImageJ was used to correct the drift between images in sequence. The coordinates of the hub assembly center were determined using the Trainable Weka Segmentation plugin (45). The MTrackJ (46) and TrackMate (47) plugins for ImageJ were used to semimanually track the coordinates of the highest pixel for each kinase domain in all CaMKII protein constructs with all HS-AFM experimental conditions. TrackMate was also used to analyze the number of detectable kinase domains. Since the diameter of the kinase domain of CaMKII is approximately 4-5 nm, we surmised that the protrusions with a diameter of approximately 4 nm around the central hub assembly consisted of the kinase domain. Using the Trackmate algorithm (Laplacian of Gaussian (LoG) detector) (*47*), the number of kinase domains in each frame of the HS-AFM videos was counted, averaged over approximately 150 frames of HS-AFM videos for each condition, and plotted.

The motions of each kinase domain of rat CaMKIIα (18,924 points in 1,577 frames of 144 kinase domains in 12 oligomers), rat nlCaMKIIα (13,740 points in 1,145 frames of 96 kinase domains in 8 oligomers), rat nlCaMKII$\alpha_{I321E}$ (22,632 points in 1,886 frames of 156 kinase domains in 13 oligomers), rat CaMKIIα + bosutinib (11,940 points in 995 frames of 84 kinase domains in 7 oligomers), rat CaMKIIα + ADP (12,384 points in 1,032 frames of 84 kinase domains in 7 oligomers), rat CaMKIIα + ATP (14,232 points in 1,186 frames of 96 kinase domains in 8 oligomers), rat CaMKIIα in EGTA/ADP after $Ca^{2+}$/CaM/ADP stimulation (13,692 points in 1,141 frames of 96 kinase domains in 8 oligomers), rat CaMKII$\alpha_{T286A}$ (18,000 points in 1,500 frames of 120 kinase domains in 10 oligomers), rat CaMKII$\alpha_{T286A}$ in EGTA/ATP after $Ca^{2+}$/CaM/ATP stimulation (17,976 points in 1,498 frames of 120 kinase domains in 10 oligomers), rat CaMKII$\alpha_{T305A/T306V}$ (18,000 points in 1,500 frames of 120 kinase domains in 10 oligomers), rat CaMKII$\alpha_{T305A/T306V}$ in EGTA/ATP after $Ca^{2+}$/CaM/ATP stimulation (18,000 points in 1,500 frames of 120 kinase domains in 10 oligomers), hydra CaMKIIα (17,592 points in 1,466 frames of 113 kinase domains in 10 oligomers), hydra CaMKIIα in EGTA/ATP after $Ca^{2+}$/CaM/ATP stimulation (17,100 points in 1,500 frames of 114 kinase domains in 10 oligomers), *C. elegans* CaMKII (17,148 points in 1,429 frames of 120 kinase domains in 10 oligomers), and *C. elegans* CaMKII in EGTA/ATP after $Ca^{2+}$/CaM/ATP stimulation (13,746 points in 1,183 frames of 93 kinase domains in 9 oligomers) were analyzed.

The motion of $Ca^{2+}$/CaM binding kinase domains in rat CaMKIIα without ATP (4,557 points of 1199 frames of 41 kinase domains in 9 oligomers), rat CaMKIIα + ATP (8,757 points of 1604 frames of 71 kinase domains in 11 oligomers), rat CaMKII$\alpha_{T286A}$ + ATP (14,234 points in 1,474 frames of 100 kinase domains in 10 oligomers), rat CaMKII$\alpha_{T305A/T306V}$ + ATP (14,214 points in 1,484 frames of 98 kinase domains in 10 oligomers), hydra CaMKIIα +ATP (12,805 points in 1,500 frames of 87 kinase domains in 10 oligomers), and *C. elegans* CaMKII + ATP (8,999 points in 1,038 frames of 62 kinase domains in 7 oligomers) were analyzed.

HS-AFM experiments were repeated at least three times independently with similar results.

Analysis of kinase domain trajectories

To quantify the trajectory of a single kinase domain, the distance from the center of the hub assembly to the kinase domains ($D_{h\text{-}k}$) was computed as follows:

$$D_{h-k} = \vec{R}_k(n) - \vec{R}_{hub}(n),$$

where $\vec{R}_k(n)$ is the kinase domain position in video frame n, and $\vec{R}_{hub}(n)$ is the center of the hub assembly position in video frame *n*.

To characterize a single kinase domain, the gyration radius, $R_g$, of the kinase domain trajectory was computed as the root-mean-square displacement of the kinase domain position from its average position,

$\langle\vec{R}\rangle$, i.e., $R_g^2 = (1/N)\sum_{n=1}^{N}\left(\vec{R}_k(n) - \langle\vec{R}\rangle\right)^2$,

where $\langle\vec{R}\rangle$ is the average position of a single kinase domain.

PCA

To visualize the coupled motions of the kinase domains, principal component analysis (PCA) was performed on their trajectories. Vectors on the kinase domains show the coupled motions along the vectors, and the lengths of the vectors indicate the degree of coupling. PCA was applied using the conventional method (*48*), that is, by diagonalizing the covariance matrix (*C*), defined as follows:

$C_{ij} = \langle(\vec{r}_i - \langle\vec{r}_i\rangle)(\vec{r}_j - \langle\vec{r}_j\rangle)\rangle$,

where $\vec{r}$ represents the x and y positions of the highest pixel for an individual kinase domain, and angular brackets represent the time average. By diagonalizing *C*, eigenvectors were computed.

Quantification and statistical analysis

All statistical analyses were performed using Igor Pro 9 software (WaveMetrics, USA) or GraphPAD Prism (GraphPad Software Inc., USA). A significance level of α = 0.05 was used for all analyses, and *p* values were adjusted for multiple comparisons where relevant. Unless otherwise noted, the normality of distributions was tested by the Shapiro–Wilk test. When it failed, the Kruskal–Wallis test and Mann–Whitney's *U* test were used to compare two groups. When several conditions were compared, one-way ANOVA and the Kruskal–Wallis test were used for the analysis of multiple groups with a single independent variable. The Dunn-Holland-Wolfe and Dunnett tests were used as follow-up tests to the Kruskal–Wallis test, where the Dunn-Holland-Wolfe test was used to compare every mean with every other mean, and the Dunnett test was used to compare every mean to a control mean. The *F* test was used to compare the variance of two groups: $p > 0.05$, not significant (N.S.); *$p < 0.05$, **$p < 0.01$, and ***$p < 0.001$ were considered statistically significant. Data are represented as the mean values ± SDs.

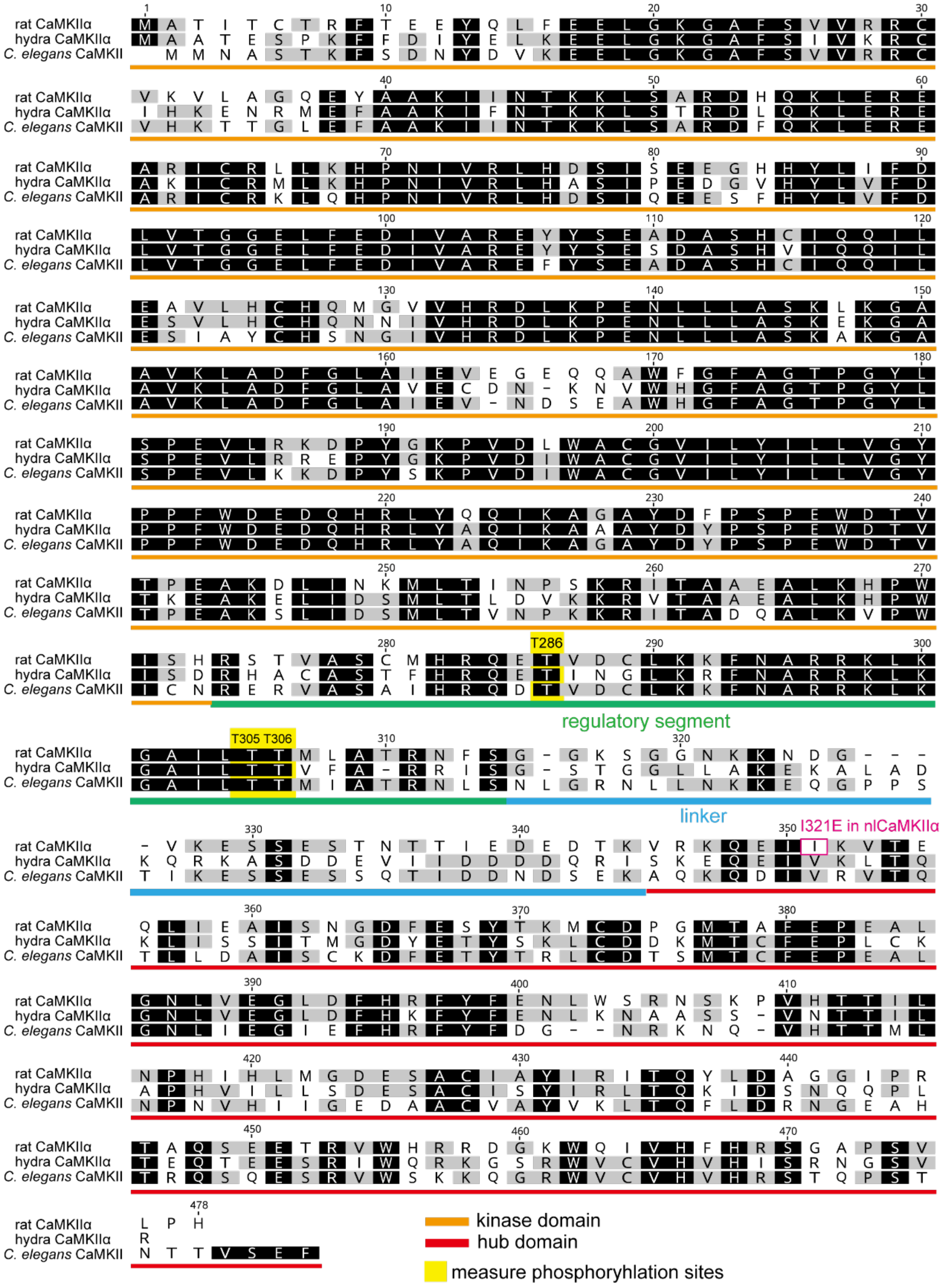


**Fig. S1. Sequence alignment of CaMKII species used in this study.**

Three CaMKII species are shown: rat CaMKIIα (*Rattus norvegicus*, Accession#: NP_037052.1), hydra CaMKIIα isoform X2 (*Hydra vulgaris*, Accession#: XP_012553992.1), and *C. elegans* CaMKII (*Caenorhabditis elegans*, Accession#: NP_001379280.1).

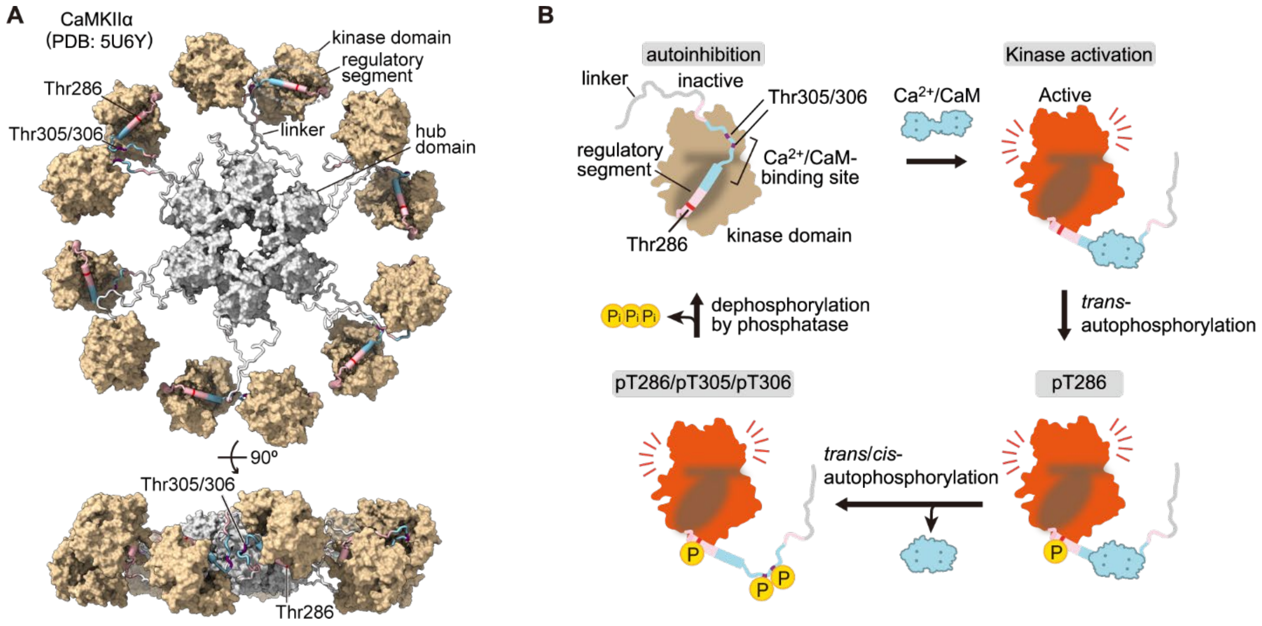


**Fig. S2. Illustration of CaMKIIα architecture.**

(**A**) Pseudoatomic model of 12-meric CaMKIIα from EM data (*12*). The solvent-excluded surface is represented. The hub domains (light gray surface, residues 345–472), kinase domains (tan surface, residues 1–273), regulatory segment (magenta cylinder, residues 274–314) with $Ca^{2+}$/CaM-binding site (blue cylinder, residues 293–310) and linker region (white ribbon, residues 315–344) are shown. Phosphorylation sites at Thr286, Thr305, and Thr306 are shown in red and purple.

(**B**) Illustration of the state change of CaMKIIα (only a single kinase domain is shown). In an inactive state, the kinase domain is autoinhibited by the binding of the regulatory segment (autoinhibition). $Ca^{2+}$/CaM binding leads to the release of the regulatory segment and the appearance of the substrate-binding site (dark tan) (i.e., kinase activation). The concomitant activation of adjacent kinases leads to autophosphorylation at Thr286 (autophosphorylation at pT286). After $Ca^{2+}$/CaM dissociation, autophosphorylation at Thr305/306 occurs. Finally, a protein phosphatase dephosphorylates CaMKIIα and returns to the inactive state.

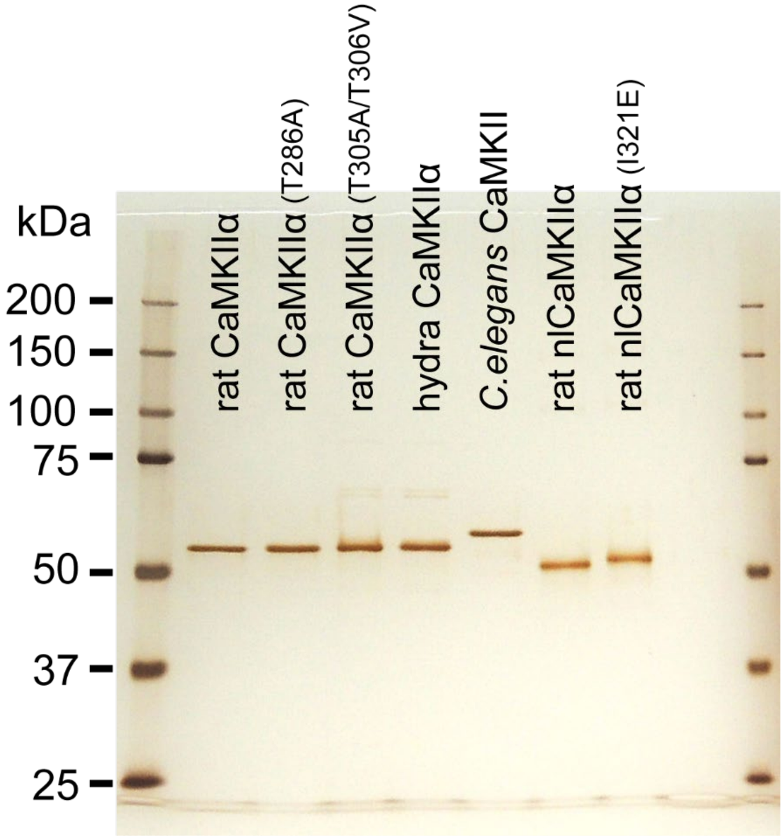


**Fig. S3. The purity of the CaMKII proteins used in this study was confirmed by silver staining.** CaMKII holoenzymes in the absence of ADP/ATP were purified from HEK-293 cells with two-step purification using His and Strep tags.

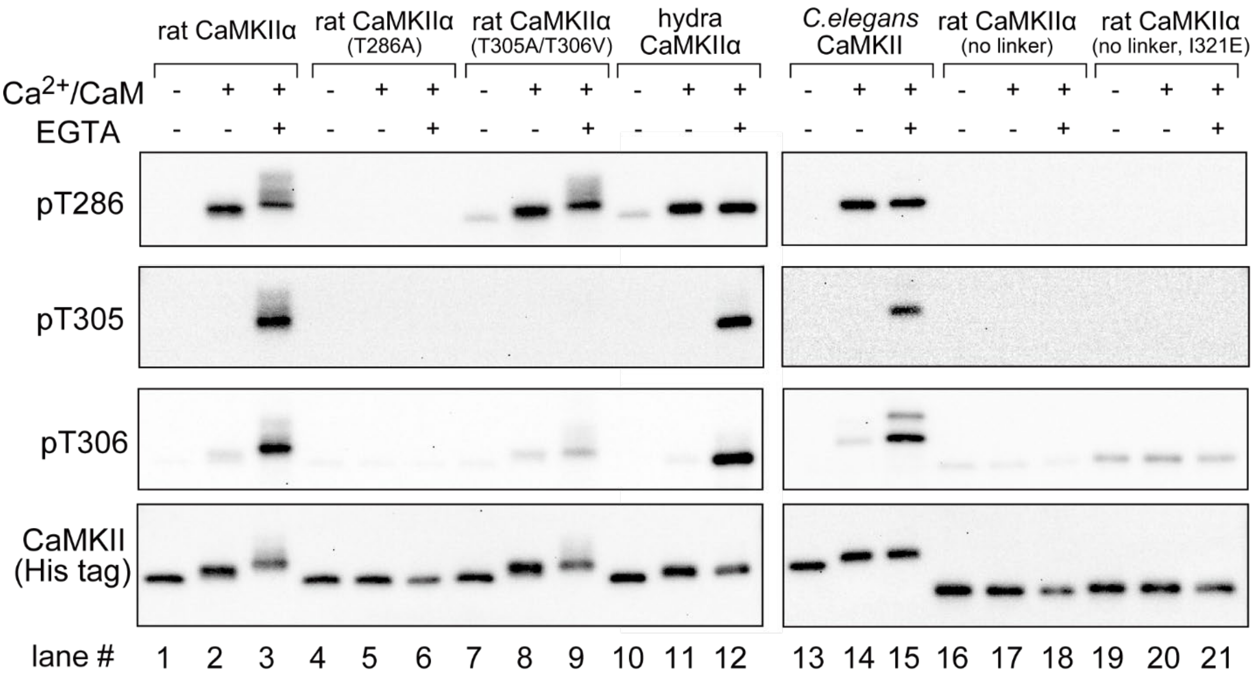


**Fig. S4. Western blotting of purified CaMKII phosphorylation.**

(lanes #1, 4, 7, 10, 13, 16, 19) Purified proteins were loaded without activation.

(lanes #2, 5, 8, 11, 14, 17, 20) CaMKII proteins (50 nM) were incubated with 800 nM CaM, 1 mM $CaCl_2$, and 1 mM ATP in a reaction buffer (50 mM Tris-HCl, pH 7.4, 150 mM KCl, and 10 mM $MgCl_2$) at 30°C for 5 min. This protocol selectively induces phosphorylation at Thr286 but not Thr305/306.

(lanes #3, 6, 9, 12, 15, 18, 21) $Ca^{2+}$/CaM was dissociated by incubation with 2 mM EGTA for 5 min at 30°C and an additional 10 min at 25°C. This protocol induces phosphorylation at Thr305/306 and kinase aggregation for rat CaMKIIα (lane #3).

The amount of protein loaded was assessed with an anti-His tag antibody.

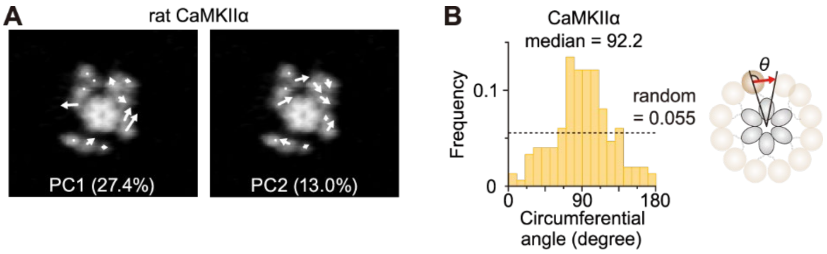


**Fig. S5. Kinase domains of the rat CaMKIIα oligomer exhibit circumferential motion around the hub assembly.**

(**A**) PCA of kinase domain motion in rat CaMKIIα. The two largest principal components of kinase domain movement are shown (PC1 and PC2). Arrows illustrate eigenvectors indicating the direction and magnitude of the collective motions of kinase domains determined by HS-AFM.

(**B**) Direction of kinase domain motions in rat CaMKIIα oligomers determined from PCA. Black dotted lines represent the random distribution.

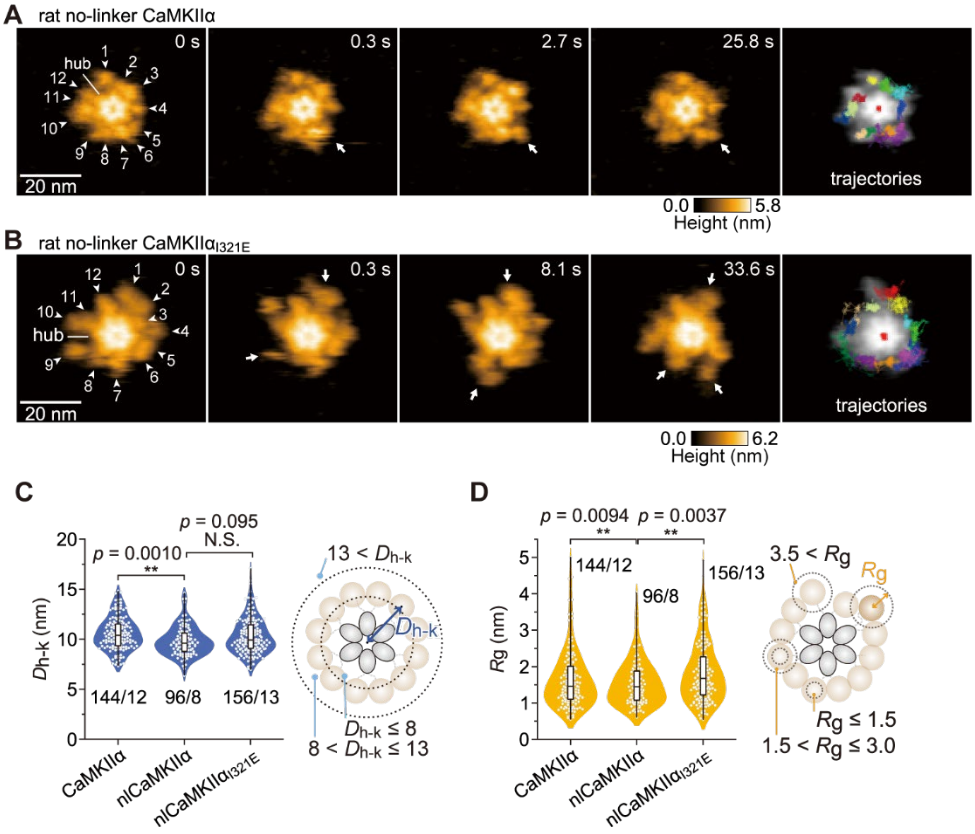


**Fig. S6. Kinase domain motion of the rat nlCaMKIIα and nlCaMKIIα$_{I321E}$ mutants.**

(**A** and **B**) Sequential HS-AFM images of rat no-linker CaMKIIα (nlCaMKIIα) (A; see also movie S1B) and rat no-linker CaMKIIα$_{I321E}$ (nlCaMKIIα$_{I321E}$) (B; see also movie S1C). White arrowheads with arbitrary numbers 1 to 12 indicate kinase domains in the oligomers. White arrows indicate the motions of kinase domains in the oligomers. Frame rate, 3.3 frames/s. Trajectories of the center of the hub assembly (red in the center) and the kinase domains in rat nlCaMKIIα (A) and rat nlCaMKIIα$_{I321E}$ (B) were tracked for ~30 s (right). Trajectories that remained roughly circular represent kinase domains that moved in a narrow range, while trajectories that covered a larger area correspond to single kinase domains that moved in a wide range.

(**C** and **D**) Distances from the center of the hub assembly to kinase domains ($D_{h\text{-}k}$) (C) and $R_g$ (D) in rat CaMKIIα, rat nlCaMKIIα, and rat nlCaMKIIα$_{I321E}$. The number of samples (kinases/holoenzymes) is indicated in the figure. N.S., not significant. $^{**}p < 0.01$ (Kruskal–Wallis test with Dunn-Holland-Wolfe post hoc test) (C). $^{**}p < 0.01$ (F test) (D). HS-AFM experiments were repeated at least three times independently with similar results.

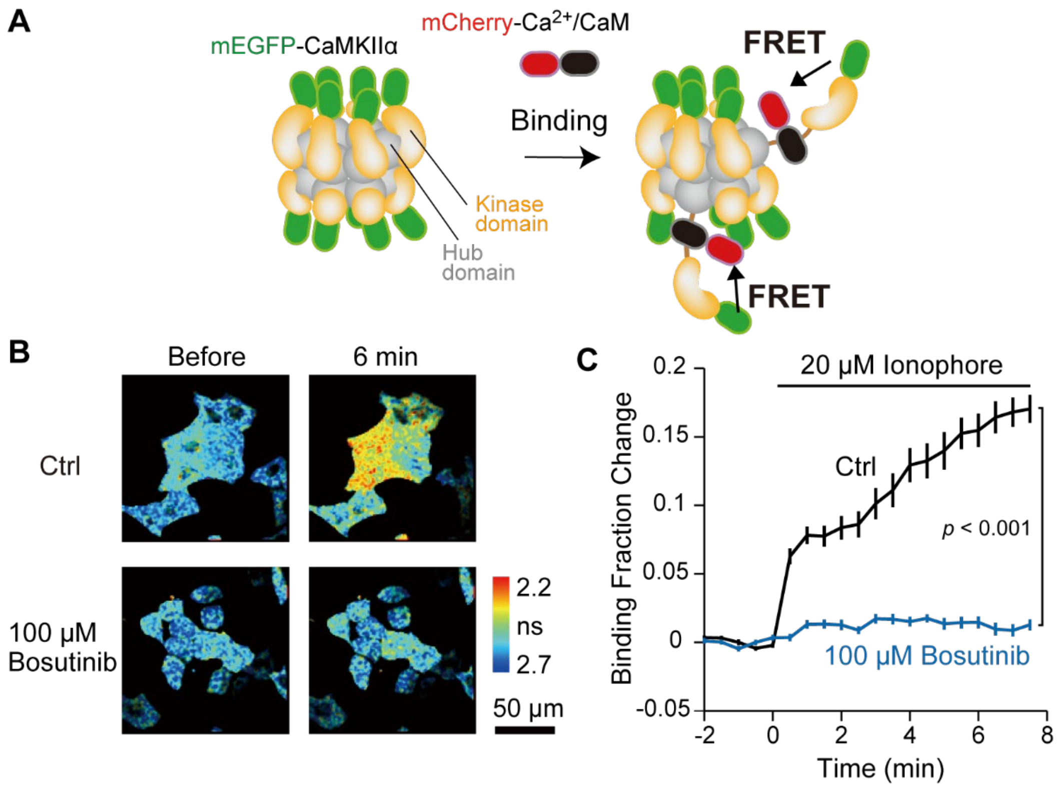


**Fig. S7. Bosutinib inhibits the binding between CaMKIIα and CaM in HeLa cells.**

(**A**) Schematic drawing of the experimental design. FRET between mEGFP-fused CaMKIIα (mEGFP-CaMKIIα) and mCherry-fused CaM was monitored by 2-photon fluorescence lifetime microscopy.

(**B**) Representative fluorescence lifetime images of HeLa cells expressing mEGFP-CaMKIIα and mCherry-CaM. The FRET donor and acceptor plasmids were transfected at a ratio of 1:3. To induce binding between mEGFP-CaMKIIα and mCherry-CaM, 20 μM ionophore was bath applied in the absence or presence of 100 μM bosutinib. Bosutinib was incubated for 40–50 min before the experiments and during the observation. The warmer color indicates the binding between mEGFP-CaMKIIα and mCherry-CaM.

(**C**) Time course of the binding fraction change after ionophore application. The number of cells analyzed was 28 for Ctrl and 33 for bosutinib. The data are shown as the mean ± sem. A t test was used for statistical comparison at 7.5 min.

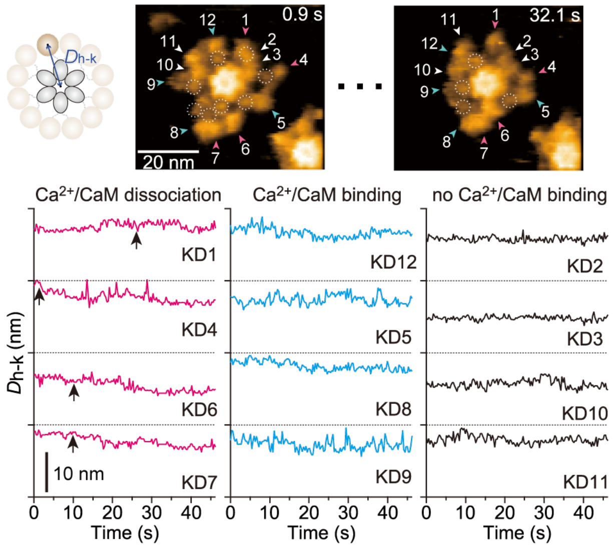


**Fig. S8. $Ca^{2+}$/CaM binding kinase domains form the extended structure in the rat CaMKIIα oligomer.**

Time course of $D_{h\text{-}k}$ of kinase domains in rat CaMKIIα (50 nM) with $Ca^{2+}$/CaM (1 mM $Ca^{2+}$, 800 nM CaM). Arbitrary numbers from 1 to 12 indicate kinase domains. White, blue, and magenta arrowheads indicate kinase domains with no binding, binding, and dissociation of $Ca^{2+}$/CaM, respectively. Black arrows in the time course indicate $Ca^{2+}$/CaM dissociation from rat CaMKIIα. KD, kinase domain. HS-AFM experiments were repeated at least three times independently with similar results.

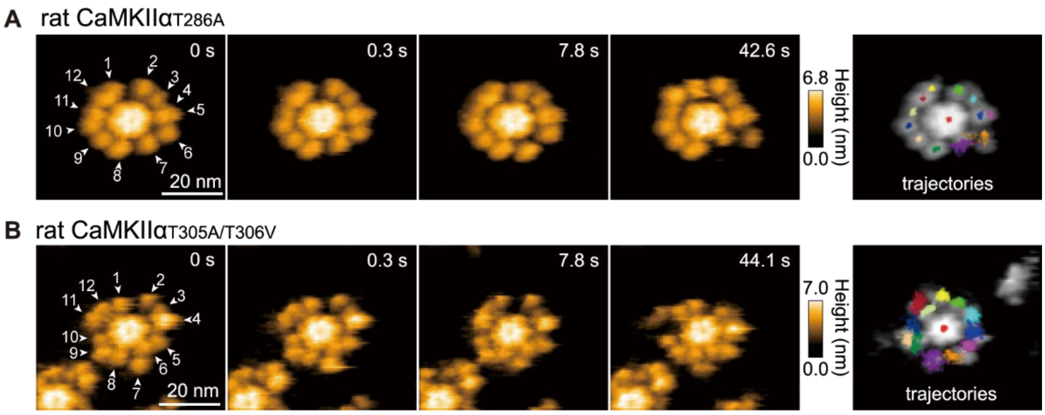


**Fig. S9. Kinase domain motion of the rat CaMKIIα$_{T286A}$ and CaMKIIα$_{T305A/T306V}$ mutants.**

(**A** and **B**) Sequential HS-AFM images of the T286A (A) and T305A/T306V (B) rat CaMKIIα mutants. White arrowheads with arbitrary numbers 1 to 12 indicate kinase domains in the oligomers. Frame rate, 3.3 frames/s. Trajectories of the center of the hub assembly (red in the center) and the kinase domains were tracked for ~30 s (right).

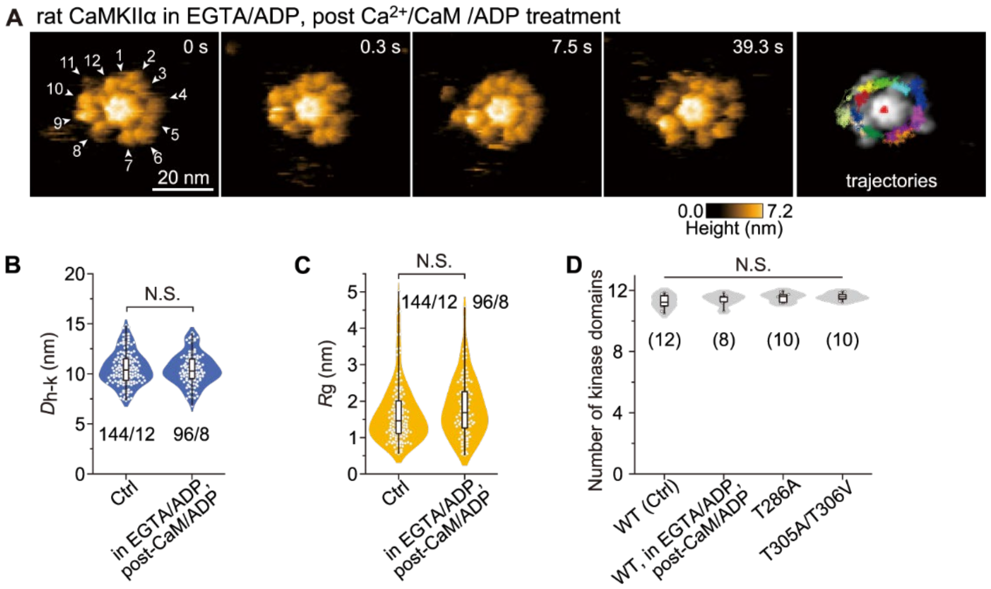


**Fig. S10. Kinase domain motion of rat CaMKIIα after $Ca^{2+}$/CaM and ADP stimulation.**

(**A**) Sequential HS-AFM images of rat CaMKIIα in EGTA (2 mM) after $Ca^{2+}$/CaM (1 mM $Ca^{2+}$, 800 nM CaM) and ADP (1 mM) stimulation. White arrowheads with arbitrary numbers from 1 to 12 indicate kinase domains in the oligomers. Frame rate, 3.3 frames/s. Trajectories of the center of the hub assembly (red in the center) and the kinase domains in rat CaMKIIα in EGTA post $Ca^{2+}$/CaM and ADP stimulation were tracked for ~30 s (right).

(**B** and **C**) Distances from the center of the hub assembly to kinase domains $D_{h\text{-}k}$ (B) and gyration radius $R_g$ (C) for rat CaMKIIα and rat CaMKIIα in EGTA after $Ca^{2+}$/CaM and ADP incubation. The number of samples (kinases/holoenzymes) is indicated in the figure. N.S., not significant (Kruskal–Wallis test).

(**D**) Number of detectable kinase domains as the 4 nm object surrounding the hub assembly in rat CaMKIIα (WT, Ctrl), WT in EGTA + ADP post $Ca^{2+}$/CaM+ADP, $CaMKII\alpha_{T286A}$, and $CaMKII\alpha_{T305/306A}$. The number of samples (holoenzymes) is indicated in the figure. No significant difference was observed (one-way ANOVA).

HS-AFM experiments were repeated at least three times independently with similar results.

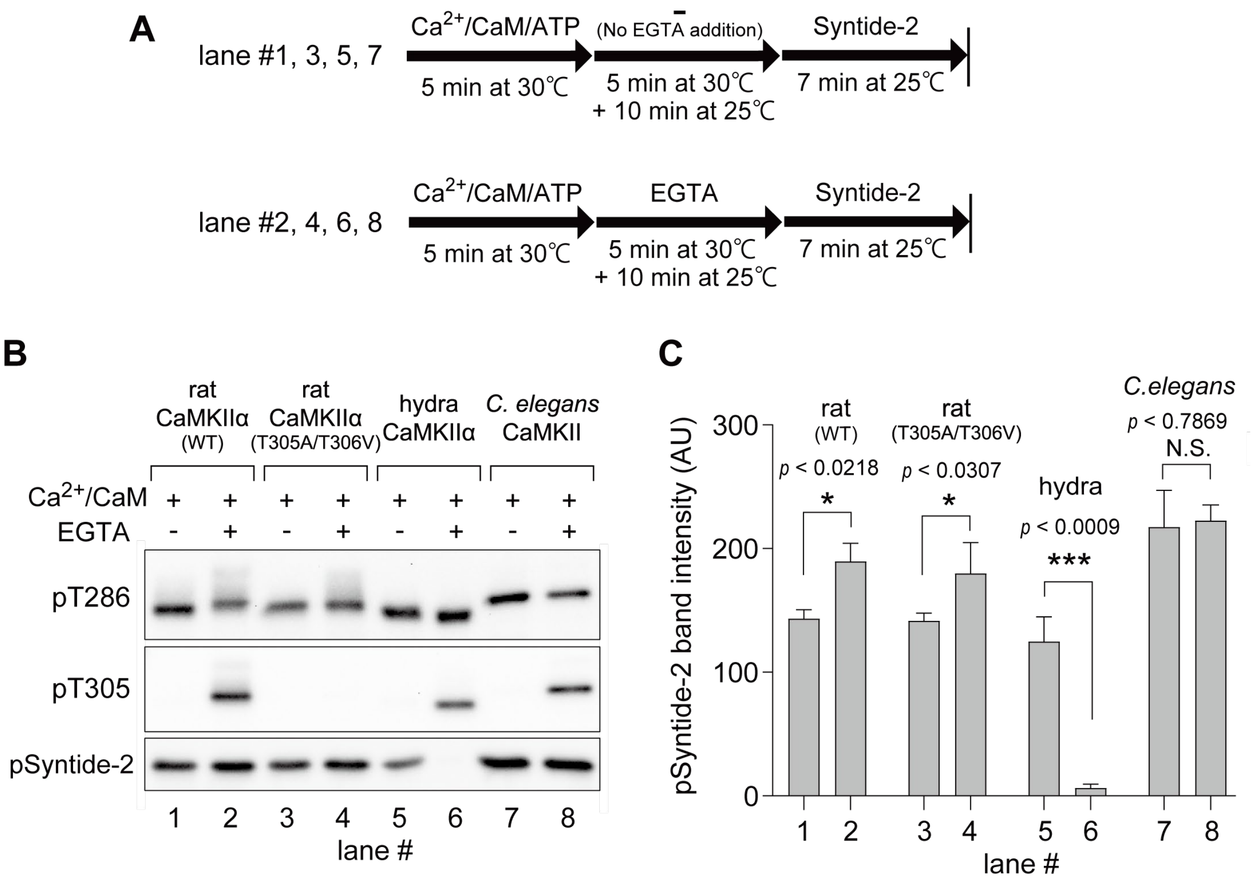


**Fig. S11. Kinase assay of pT286/pT305/pT306 CaMKII holoenzymes.**

(**A** and **B**) Phosphorylation of the sfGFP-syntide-2 peptide by pT286/pT305/pT306 CaMKII species was detected by western blotting. pT286 CaMKII (lanes 1, 3, 5, 7) and pT286/pT305/pT306 CaMKII (lanes 2, 4, 6, 8) were prepared as described in the Materials and Methods. This protocol also induces kinase aggregation in rat CaMKIIα holoenzymes (Fig. 3B). Subsequently, 1 μM sfGFP-Syntide-2 was incubated for 7 min at 25°C as a substrate for CaMKII.

(**C**) Quantification of the experiments. For data analysis, kinase activity. Error bars indicate SEM for four independent experiments. N.S. (not significant, $p > 0.05$); *$p < 0.05$; ***$p < 0.001$; paired *t* test.

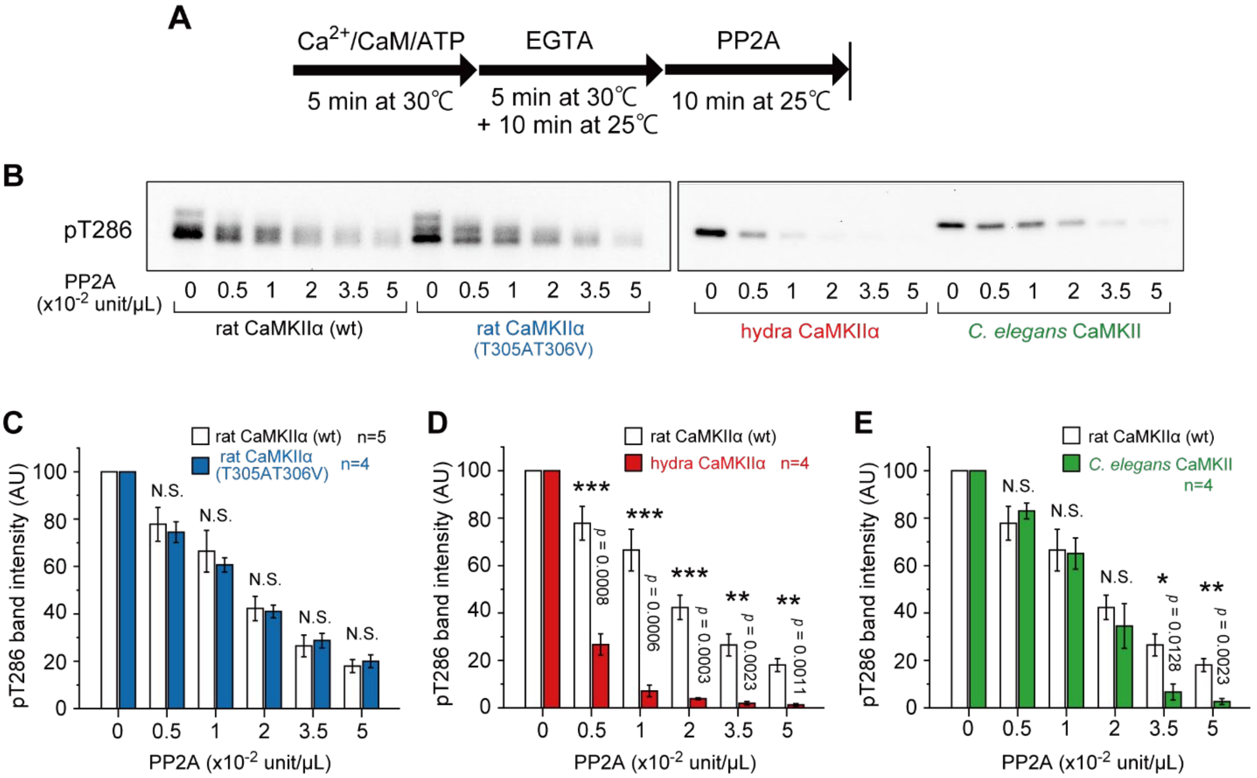


**Fig. S12. Assay of pT286/pT305/pT306 CaMKII holoenzyme dephosphorylation.**

(**A** and **B**) Dephosphorylation of CaMKII by PP2A was detected by western blotting. The time course of the experiment (A) and the representative result of western blotting (B) are shown. pT286/pT305/pT306 CaMKII was prepared as described in the Materials and Methods. This protocol also induces kinase aggregation in rat CaMKIIα holoenzymes (Fig. 3B). Subsequently, the indicated amount of PP2A was incubated for 10 min at 25°C.

(**C** to **E**) Statistical comparison of rat CaMKIIα (wt) and the T305A/T306V mutant (C), hydra CaMKIIα (D), and *C. elegans* CaMKII (E). Error bars indicate SEM, and the number of experiments is indicated in the figures. N.S. (not significant, $p > 0.05$); *$p < 0.05$; ***$p < 0.001$; *t* test.

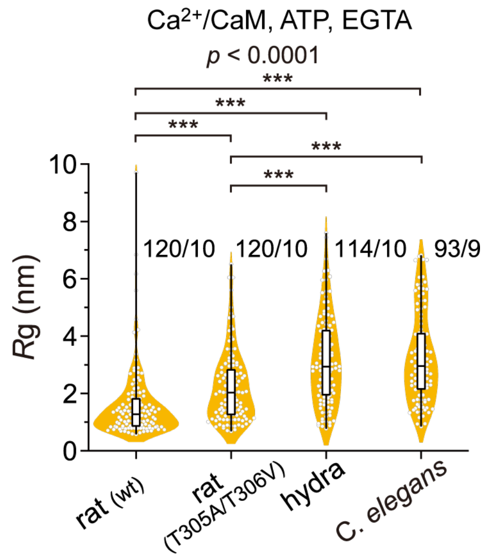


**Fig. S13. Statistical comparison of $R_g$ of the fully phosphorylated CaMKII from different species.**

The same data set used in Fig. 3G and Fig. 4H was used here. $^{*}p < 0.05$, $^{***}p < 0.001$ (Kruskal–Wallis test with Dunn's post hoc test). The number of samples (kinases/holoenzymes) is indicated in the figure.

**Movie S1. HS-AFM videos of three representative rat CaMKIIα$_{WT}$ and no-linker CaMKIIα molecules on a P[5]A+-modified mica surface. (A)** Rat CaMKIIα$_{WT}$ (50 nM). (**B**) Rat no-linker CaMKIIα (nlCaMKIIα). (**C**) Rat no-linker CaMKIIα$_{I321E}$ (nlCaMKIIα$_{I321E}$). The trajectories of the kinase domains in the rat CaMKIIα oligomers are also shown at the bottom. Image size, 110 × 98 pixels$^2$; scan area, 62 × 55 nm$^2$; frame rate, 3.3 fps.

**Movie S2. HS-AFM videos of three representative rat CaMKIIα$_{WT}$ molecules on a P[5]A+-modified mica surface.** (**A**) Rat CaMKIIα$_{WT}$ (50 nM) without any addition of inhibitors or ligands (ctrl). These HS-AFM videos are the same as Movie S1A. (**B, C**) Rat CaMKIIα$_{WT}$ treated with 50 μM bosutinib (B) and $Ca^{2+}$/CaM (1 mM $Ca^{2+}$, 800 nM CaM (C). The trajectories of the kinase domains in the CaMKIIα oligomers are also shown at the bottom. Image size, 110 × 98 pixels$^2$; scan area, 62 × 55 nm$^2$; frame rate, 3.3 fps.

**Movie S3. HS-AFM videos of three representative rat CaMKIIα$_{WT}$ molecules after phosphorylation on a P[5]A+-modified mica surface.** (**A**) Rat CaMKIIα$_{WT}$ (50 nM) (ctrl). These HS-AFM videos are the same as MovieS1A. (**B, C**) Rat CaMKIIα$_{WT}$ treated with $Ca^{2+}$/CaM + ATP (1 mM $Ca^{2+}$, 800 nM CaM, 1 mM ATP) (B) and EGTA (2 mM) + ATP (1 mM) after $Ca^{2+}$/CaM/ATP stimulation (1 mM $Ca^{2+}$, 800 nM CaM, 1 mM ATP) (C). The trajectories of the kinase domains in the CaMKIIα oligomers are also shown at the bottom. Dotted white circles indicate $Ca^{2+}$/CaMs bound to CaMKIIα during the first 9.9 s. Image size, 110 × 98 pixels$^2$; scan area, 62 × 55 nm$^2$; frame rate, 3.3 fps.

**Movie S4. HS-AFM videos of three representative rat CaMKIIα$_{T286A}$ molecules on a P[5]A+-modified mica surface.** (**A**) Rat CaMKIIα$_{T286A}$ (50 nM) (ctrl). (**B, C**) Rat CaMKIIα$_{T286A}$ treated with $Ca^{2+}$/CaM + ATP (1 mM $Ca^{2+}$, 800 nM CaM, 1 mM ATP) (B) and EGTA (2 mM) + ATP (1 mM) after $Ca^{2+}$/CaM/ATP stimulation (1 mM $Ca^{2+}$, 800 nM CaM, 1 mM ATP) (C). The trajectories of the kinase domains in the CaMKIIα oligomers are also shown at the bottom. Dotted white circles indicate $Ca^{2+}$/CaMs bound to CaMKIIα during the first 9.9 s. Image size, 110 × 98 pixels$^2$; scan area, 62 × 55 nm$^2$; frame rate, 3.3 fps.

**Movie S5. HS-AFM videos of three representative rat CaMKIIα$_{T305A/T306V}$ molecules on a P[5]A+-modified mica surface.** (**A**) Rat CaMKIIα$_{T305A/T306V}$ (50 nM) (ctrl). (**B, C**) Rat CaMKIIα$_{T305A/T306V}$ treated with $Ca^{2+}$/CaM + ATP (1 mM $Ca^{2+}$, 800 nM CaM, 1 mM ATP) (B) and in EGTA (2 mM) + ATP (1 mM) after $Ca^{2+}$/CaM/ATP stimulation (1 mM $Ca^{2+}$, 800 nM CaM, 1 mM ATP) (C). The trajectories of the kinase domains in the CaMKIIα oligomers are also shown at the bottom. Dotted white circles indicate $Ca^{2+}$/CaMs bound to CaMKIIα during the first 9.9 s. Image size, 110 × 98 pixels$^2$; scan area, 62 × 55 nm$^2$; frame rate, 3.3 fps.

**Movie S6. HS-AFM videos of three representative hydra CaMKIIα$_{WT}$ molecules on a P[5]A+-modified mica surface.** (**A**) hydra CaMKIIα$_{WT}$ (50 nM) (ctrl). (**B, C**) Hydra CaMKIIα$_{WT}$ treated with $Ca^{2+}$/CaM + ATP (1 mM $Ca^{2+}$, 800 nM CaM, 1 mM ATP) (B) and EGTA (2 mM) + ATP (1 mM) after $Ca^{2+}$/CaM/ATP stimulation (1 mM $Ca^{2+}$, 800 nM CaM, 1 mM ATP) (C). The trajectories of the kinase domains in the CaMKIIα oligomers are also shown at the bottom. Dotted white circles indicate $Ca^{2+}$/CaMs bound to CaMKIIα during the first 9.9 s. Image size, 110 × 98 pixels$^2$; scan area, 62 × 55 nm$^2$; frame rate, 3.3 fps.

**Movie S7. HS-AFM videos of three representative *C. elegans* CaMKII$_{WT}$ molecules on a P[5]A+-modified mica surface.** (**A**) *C. elegans* CaMKII$_{WT}$ (50 nM) (ctrl). (**B, C**) *C. elegans* CaMKII$_{WT}$ treated with $Ca^{2+}$/CaM + ATP (1 mM $Ca^{2+}$, 800 nM CaM, 1 mM ATP) (B) and EGTA (2 mM) + ATP (1 mM) after $Ca^{2+}$/CaM/ATP stimulation (1 mM $Ca^{2+}$, 800 nM CaM, 1 mM ATP) (C). The trajectories of the kinase domains in the CaMKIIα oligomers are also shown at the bottom. Dotted white circles indicate $Ca^{2+}$/CaMs bound to CaMKIIα during the first 9.9 s. Image size, 110 × 98 pixels$^2$; scan area, 62 × 55 nm$^2$; frame rate, 3.3 fps.
